# Immunoresolvents Support Skeletal Myofiber Regeneration via Actions on Myeloid and Muscle Stem Cells

**DOI:** 10.1101/2020.06.12.148320

**Authors:** James F. Markworth, Lemuel A. Brown, Eunice Lim, Carolyn Floyd, Jaqueline Larouche, Jesus A. Castor-Macias, Kristoffer B. Sugg, Dylan C. Sarver, Peter C. D. Macpherson, Carol Davis, Carlos A. Aguilar, Krishna Rao Maddipati, Susan V. Brooks

**Affiliations:** Department of Molecular & Integrative Physiology, University of Michigan, Ann Arbor, Michigan; Department of Orthopaedic Surgery, University of Michigan, Ann Arbor, Michigan; Department of Surgery, University of Michigan, Ann Arbor, Michigan; Department of Cellular & Molecular Physiology, Johns Hopkins University, Baltimore, Maryland; Department of Biomedical Engineering, University of Michigan, Ann Arbor, Michigan; Department of Pathology, Lipidomics Core Facility, Wayne State University, Detroit, Michigan

## Abstract

Specialized pro-resolving mediators (SPMs) actively limit inflammation and expedite its resolution. Here we profiled intramuscular lipid mediators following injury and investigated the role of SPMs in skeletal muscle inflammation and repair. Both eicosanoids and SPMs increased following myofiber damage induced by intramuscular injection of barium chloride or functional overload. Daily systemic administration of resolvin D1 (RvD1) limited the degree and duration of inflammation, enhanced regenerating myofiber growth, and improved recovery of muscle strength. RvD1 suppressed inflammatory cytokines, enhanced polymorphonuclear cell clearance, modulated muscle stem cells, and polarized macrophages to a more pro-regenerative subset. RvD1 had minimal direct impact on *in-vitro* myogenesis but directly suppressed myokine production and stimulated macrophage phagocytosis, showing that SPMs influence modulate both infiltrating myeloid and resident muscle cells. These data reveal the efficacy of immunoresolvents as a novel alternative to classical anti-inflammatory interventions in the management of muscle injuries to modulate inflammation while stimulating tissue repair.

## Introduction

Skeletal muscle damage induces an acute inflammatory response, during which blood leukocytes are recruited to the site of injury (1). Polymorphonuclear cells (PMNs) first appear within muscle in the early hours to days following injury (2). The initial wave of PMNs is followed by recruitment of blood monocytes that locally differentiate into tissue macrophages (MΦ) (3). The MΦ that appear initially following muscle injury exhibit a phagocytic pro-inflammatory phenotype, invading within necrotic myofibers to clear tissue debris (4). These phagocytic MΦ also likely play an important role in the uptake and removal of apoptotic cells such as PMNs within the injured musculature (5). In addition to preparing the injury site to enable subsequent repair, phagocytosis acts as a stimulatory cue to trigger a switch to a reparative MΦ subset, which supports muscle regeneration via interactions with post-mitotic muscle cells (myofibers) and associated resident myogenic stem cells (muscle satellite cells, MuSCs) (6). Unlike MΦ, current evidence suggests that PMNs have overall deleterious effects on muscle regeneration (7). Therefore, it is important that PMN influx ceases in a timely manner and that the PMNs which invaded muscle initially following injury are cleared from regenerating tissue, such that the inflammatory response is successfully resolved (5).

Resolution of the acute inflammatory response was classically considered to be a passive event resulting from dilution and removal of pro-inflammatory mediators produced following injury (8). More recently, it has been demonstrated that distinct families of specialized pro-resolving lipid mediators (SPMs) are produced during the resolution phase of the acute inflammatory response (9), via the coordinated action of the 5-, 12- and 15-lipoxygenase (LOX) pathways (10). SPMs are a superfamily of bioactive metabolites of dietary essential fatty acids including the lipoxins [derived from omega-6 (n-6) arachidonic acid (20:4n-6)] (11), the E-series resolvins [derived from n-3 eicosapentaenoic acid (EPA; 20:5n-3)] (12), and the D-series resolvins (13), protectins (14), and maresins (15) [all derived from n-3 docosahexaenoic acid (DHA; 22:6n-3)]. These autocoids actively limit further PMN influx, while simultaneously stimulating MΦ functions essential for timely resolution of inflammation and tissue repair, such as enhancing their phagocytic capacity and promoting a M2-like polarization state (16). Emerging evidence suggests that SPMs can also exert important biological actions on various non-immune cell populations (e.g. stem cells) that may contribute to their physiological effects (17).

The role of lipid mediators in skeletal muscle inflammation and repair is poorly understood (18). Prior studies have largely focused on the prostaglandins, classical pro-inflammatory eicosanoid metabolites of n-6 arachidonic acid synthesized via the cyclooxygenase (COX)-1 and 2 pathways (19). Given the importance of inflammation in the control of muscle remodeling, it is not surprising that many studies have found that systemic treatment with non-steroidal anti-inflammatory drugs (NSAIDs) can impede skeletal muscle regeneration (20–25). The discovery of SPMs has allowed for development of novel strategies to modulate inflammation which differ fundamentally from this classical anti-inflammatory approach (26). Administration of resolution agonists, coined immunoresolvents, such as native SPMs or drug mimetics, can limit inflammation and expedite its resolution, while simultaneously relieving pain in many clinically relevant models of acute and chronic inflammation (27). Unlike NSAIDs, SPMs have been recently found to have pro-regenerative tissue actions following dermal wound healing (28–30), spinal cord injury (31), bone fracture (32, 33), and myocardial infarction (34–37).

SPMs are locally produced and appear systemically following skeletal muscle injury in both mice and humans, suggesting that they may play an important role in control of the inflammatory and adaptive responses to myofiber damage (18, 38–42). Recently, intramuscular injection to MuSC deficient mice of one native SPM, resolvin D2 (RvD2), was reported to accelerate MΦ transitions following degenerative muscle injury, resulting in improved recovery of muscle mass and strength (38). However, the endogenous role of SPMs and the therapeutic applicability of immunoresolvents to modulate muscle inflammation and repair under physiological circumstances where MuSCs play an indispensable role in successful myofiber regeneration remains unknown (43). Moreover, this prior study focused on immunological endpoints, thus there remains a complete lack of data concerning the impact of immunoresolvents on cellular and molecular indices of myofiber growth/regeneration or the MuSC response (38). Recently, Annexin A1, a key pro-resolving peptide which shares its cell surface receptor (FPR2/ALX) with the SPM resolvin D1 (RvD1), was also shown to control muscle regeneration in healthy young mice by muscle cell direct and indirect actions (44).

In the current study, we investigated the potential role of SPMs in modulating inflammatory and regenerative responses in physiological models of skeletal muscle growth and repair including both adaptive myofiber hypertrophy and myofiber degeneration/regeneration. Moreover, we tested whether daily systemic administration of RvD1, as an exogenous immunoresolvent, regulates cellular and molecular indices of myofiber regeneration and myogenic MuSC responses to muscle injury in healthy young mice *in vivo* and whether RvD1 directly modulates myogenesis and myokine production *in vitro*.

## Results

### Temporal Changes in Muscle Lipid Mediators Following Muscle Injury

Intramuscular injection of barium chloride (BaCl_2_) resulted in widespread myofiber damage and local appearance of both PMN (Ly6G^+^ cells) and MΦ (CD68^+^ cells) (Figure 1A). By day 3, there was progressive PMN clearance and a robust infiltration of MΦ. Many small regenerating myofibers with characteristic centrally located nuclei appeared by day 5, together with a persistent MΦ presence. Muscle COX-2 expression increased markedly at day 1 post-injury, but intramuscular lipid mediator profile as determined by LC-MS did not yet differ substantially from sham injured mice (Figure 1B-C). By day 3, there were increased pro-inflammatory eicosanoids derived from the COX (e.g. TXB_2_, PGE_2_ and PGD_2_) and 12-LOX pathways (e.g. 12-HETE). Muscle expression of 5-lipoxygenase (5-LOX), platelet-type 12-lipoxygenase (12S-LOX), and leukocyte-type 12/15-lipoxygenase (12/15-LOX) increased later, resulting in increased concentrations of LOX generated monohydroxylated pathway intermediates in the biosynthesis of pro-inflammatory leukotrienes (5-HETE), as well as SPMs including the lipoxins (15-HETE), E-series resolvins (18-HEPE), D-series resolvins/protectins (17-HDoHE), and maresins (14-HDoHE) (Figure 1C). Downstream bioactive SPMs including protectin D1 and maresin 1 were also detected at day 3 post-injury, although lipoxins, E-resolvins, and D-resolvins were generally below the limits of detection within muscle (Supplemental Tables 1a). In addition, we detected for the first time elevated levels of many metabolites of epoxygenases (CYP p450) in injured mouse muscle (Figure 1C, Supplemental Tables 1b).

**Figure 1.**
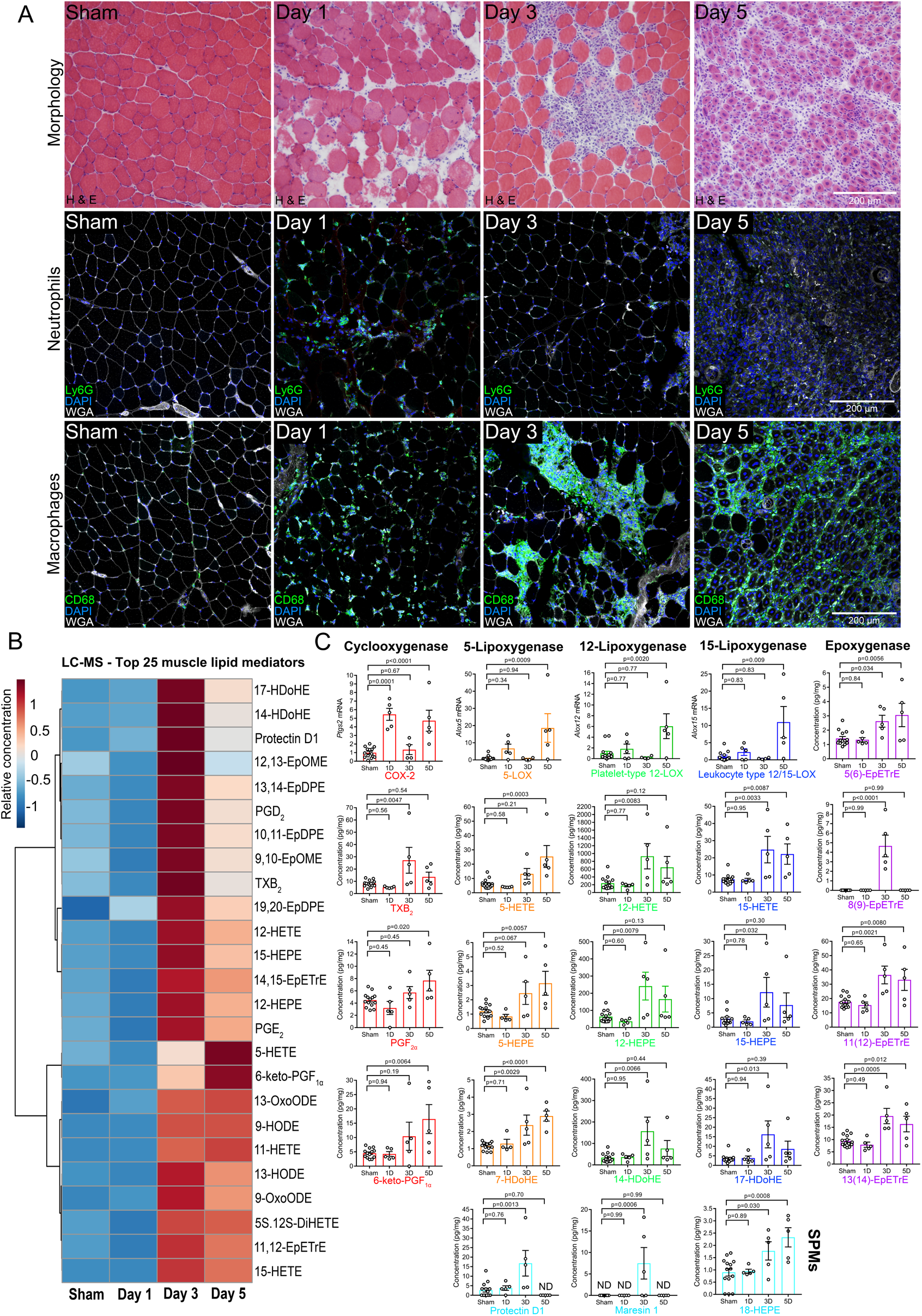
Dynamic shifts in muscle lipid mediators following skeletal muscle injury: A: Female C57BL/6 mice (4-6 mo) received bilateral intramuscular injection of the tibialis anterior (TA) muscle with 50 μL of 1.2% barium chloride (BaCl_2_) to induce myofiber injury. Age and gender matched mice received bilateral sham intramuscular injections with 50 μL of sterile saline. TA cross-sections were stained for hematoxylin and eosin (H&E), polymorphonuclear cells (PMNs, Ly6G), or monocytes/macrophages (MΦ, CD68). Scale bars are 200 μm. B: Heat map of the top 25 muscle lipid mediators (based on PLS-DA VIP score) modulated by muscle injury as determined by LC-MS analysis. C: Whole muscle mRNA expression of major murine lipid mediator biosynthetic enzymes and respective downstream lipid metabolites as measured by LC-MS including the cyclooxygenase-2 (COX-2, red), 5-lipoxygenase (5-LOX, orange), platelet-type 12-lipoxygenase (12-LOX, green), leukocyte-type 12/15-lipoxygenase (12/15-LOX, dark blue) and epoxygenase (CYP 450, purple) pathways. Downstream bioactive specialized pro-resolving lipid mediators (SPMs) detected in muscle are also shown (light blue). Bars show the mean ± SEM of 4-5 mice per group with dots representing data from each individual mouse (biological replicates).The sham injury time-course was pooled for quantitative analysis and is shown in full in Supplemental Tables 1c. P-values were determined by one-way ANOVA followed by Holm-Sidak post-hoc tests with the pooled sham injury group serving as control. ND denotes analytes that were below the limits of detection.

### Dynamic Changes in Muscle Lipid Mediators Are Conserved Across Different Species and Models of Injury

We also performed LC-MS based lipidomic profiling of the plantaris muscle following synergist ablation-induced functional overload in rats (Figure 2A). This is a milder, but potentially more physiologically relevant model of myofiber damage when compared to BaCl_2_ induced injury (45) (Figure 2A). Synergist ablation resulted in a compensatory increase in the mass of the overloaded plantaris at 28 days’ post-surgery (Figure 2B-C). This was attributable to an increase in muscle fiber size (Figure 2D), and was most evident for slow-twitch Type I and IIa fiber types (46) (Figure 2E). Control plantaris muscles contained many resident ED2 (CD163^+^) MΦ, few scattered ED1 (CD68^+^) MΦ, and very few PMNs (HIS-48^+^ cells). Therefore, unlike in mice, the resident MΦ in rat muscle were predominantly CD163 (ED2^+^), but CD68 (ED1^−^) (47) (Supplemental Figure 1). Synergist ablation resulted in localized inflammation of the overloaded plantaris at day 3 post-surgery (Figure 2A, Supplemental Figure 2A). At least three distinct myeloid cell populations were present including PMNs (HIS-48^+^CD68^−^CD163^−^ cells), inflammatory ED1 MΦ (HIS-48^−^CD68^+^CD163^−^ cells), and resident-like ED2 MΦ (HIS-48^−^CD68^−^CD163^+^ cells) (Figure 2F). Scattered HIS48^+^ cells could still be seen at day 7, but were absent by day 28. In contrast, CD68^+^ and CD163^+^ cells persisted, albeit in much lower numbers, at both 7 and 28 days of recovery, with a clear increase in co-expression of CD68 and CD163 by the MΦ which remained (Supplemental Figure 2B).

**Figure 2.**
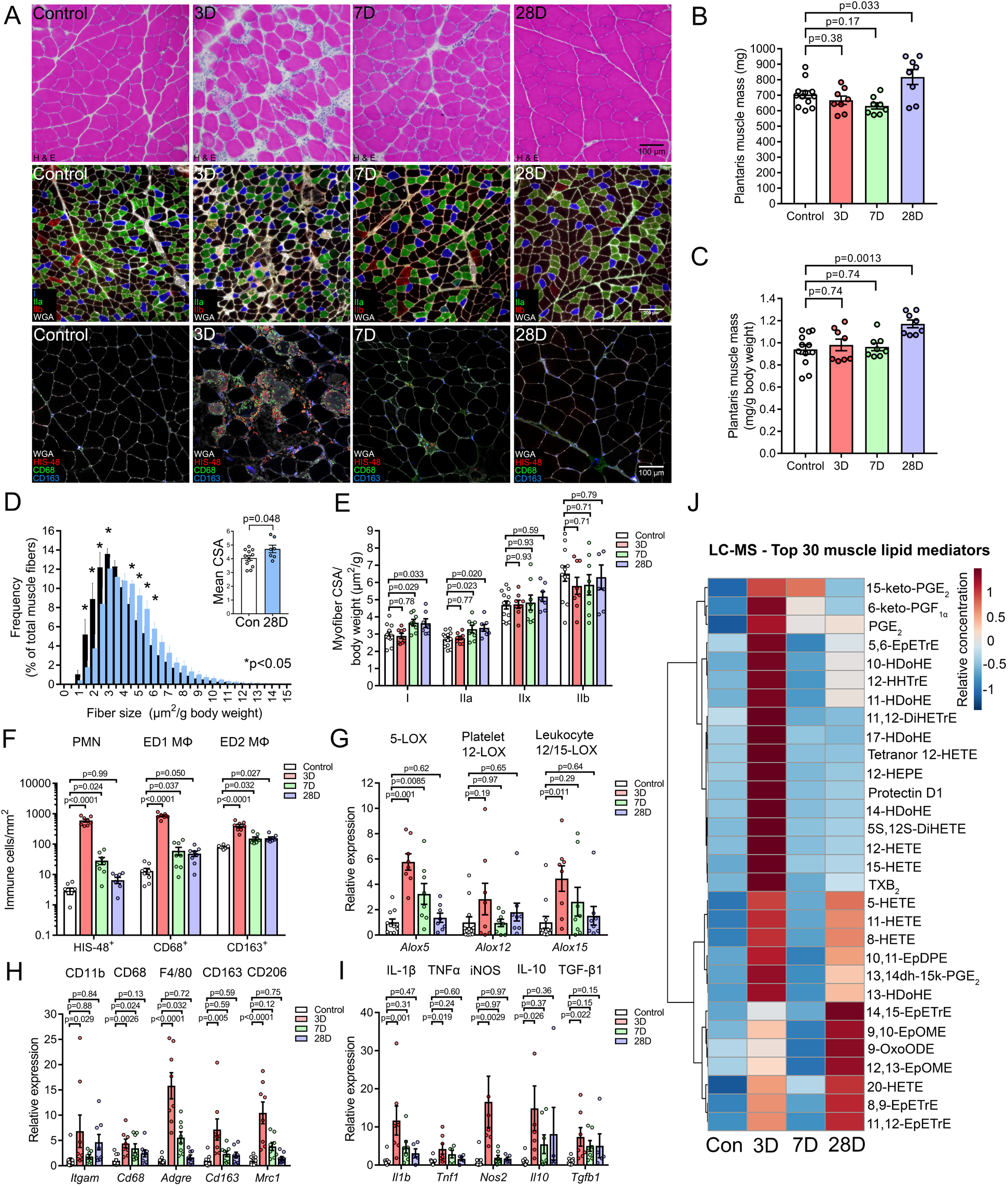
Muscle lipid mediator responses to functional plantaris overload: A: Male Sprague Dawley rats (6 mo) underwent bilateral functional overload of the plantaris muscle via synergist ablation surgery. Plantaris muscles from age and gender matched rats served as non-surgical controls. Muscle cross-sections were stained for hematoxylin & eosin (H&E), muscle fiber type, and inflammatory cells including polymorphonuclear cells (PMNs, HIS-48), ED1 monocyte/macrophages (MΦ, CD68), and ED2 MΦ (CD163). Scale bars are 100 μm. B: Absolute plantaris muscle mass (mg) following functional overload C: Relative plantaris muscle mass (mg/g body weight) following functional overload. D: Percentage frequency distribution of cross sectional area (CSA) of total muscle fiber population in plantaris muscles of control and day 28 post-synergist ablation rats. The inset shows the mean myofiber CSA. E: Mean muscle fiber CSA of myofibers in the overloaded plantaris muscle split by respective muscle fiber type. F: Quantification of intramuscular PMNs (HIS-48^+^ cells), inflammatory ED1 MΦ (CD68^+^ cells), and resident/M2-like ED2 MΦ (CD163^+^ cells). G: Whole muscle mRNA expression of 5-lipoxygenase (5-LOX), platelet-type 12-lipoxygenase (12-LOX), and leukocyte type 12/15-lipoxygenase (12/15-LOX). H-I: Whole muscle mRNA expression of immune cell markers, cytokines, and markers of MΦ activation state. Gene of interest expression was normalized to Gapdh. J: Heat map of the top 30 muscle lipid mediators (based on PLS-DA VIP score) modulated by functional overload as determined by LC-MS analysis. The full LC-MS data set is shown in Supplemental Tables 1d. Bars show the mean ± SEM of 8-12 plantaris muscles from 4-6 rats per group, with each surgical site (limb) considered to be a biological replicate. Dots represent data from each individual muscle sample. P-values were determined one-way ANOVA followed by Holm-Sidak post-hoc tests with non-surgery rats serving as a control group (panels B-C and E-I) or by two-tailed unpaired t-tests (panel D).

Plantaris muscle inflammation was accompanied by increased muscle mRNA expression of major rat 5-, 12-, and 15-LOX enzymes (Figure 2G), immune cell markers (Figure 2H), and cytokines/MΦ activation markers (Figure 2I) (Table 2). LC-MS based lipidomic profiling showed elevated intramuscular concentrations of lipid mediators from the COX, LOX, and CYP pathways (Figure 2J, Supplemental Tables 1e). These included pro-inflammatory eicosanoids (e.g. PGE_2_, TXB_2_, 5-HETE, 12-HETE), as well as pathway markers of SPM biosynthesis including the lipoxins (15-HETE), D-series resolvins/protectins (17-HDoHE), and maresins (14-HDoHE). Bioactive SPMs including lipoxin A_4_, protectin D1, maresin 1, resolvin D6, and 8-oxo-Resolvin D1 were also detected (Supplemental Tables 1f). While most COX and LOX metabolites had returned to resting levels by day 7, many CYP pathway metabolites increased later at day 28 (Figure 2J, Supplemental Tables 1g).

### Systemic Resolvin D1 Treatment Limits Muscle Inflammation

We next investigated the role of SPMs in controlling acute muscle inflammation. For this, we returned to the mouse BaCl_2_ model in which muscle inflammation was more uniform and wide-spread. Mice were treated with resolvin D1 (RvD1), a native SPM derived endogenous from n-3 DHA by the sequential actions of the 15- and 5-LOX pathways (48, 49). We chose to test RvD1 as a prototypical immunoresolvent due to its extensively documented receptor dependent pro-resolving actions (50–55), as well as its established *in-vivo* pharmacokinetics and previously demonstrated therapeutic efficacy following systemic administration in mice (56–60). Intraperitoneal (IP) injection of RvD1 at the time of BaCl_2_ injury reduced intramuscular MΦ (CD68^+^ cells) 24 h later, but did not affect PMN (Ly6G^+^ cell) number at this time-point (Figure 3A). RvD1 treatment also reduced muscle mRNA expression of the pan innate immune cell marker CD11b, MΦ markers including CD68, F4/80, and CD206 (Figure 3B), as well as cytokines including IL-6, IL-1β, MCP-1, and TNFα (Figure 3B). Common housekeeping genes including both beta-actin and 18S were markedly induced together with genes of interest in response to intramuscular BaCl_2_ injection and RvD1 treatment reduced 18S, but not beta-actin expression (data not shown). Therefore, RT-PCR data in Figure 3 was normalized to beta-actin (*Actb*) expression and presented relative to injured vehicle treated mice for quantitative analysis.

**Figure 3.**
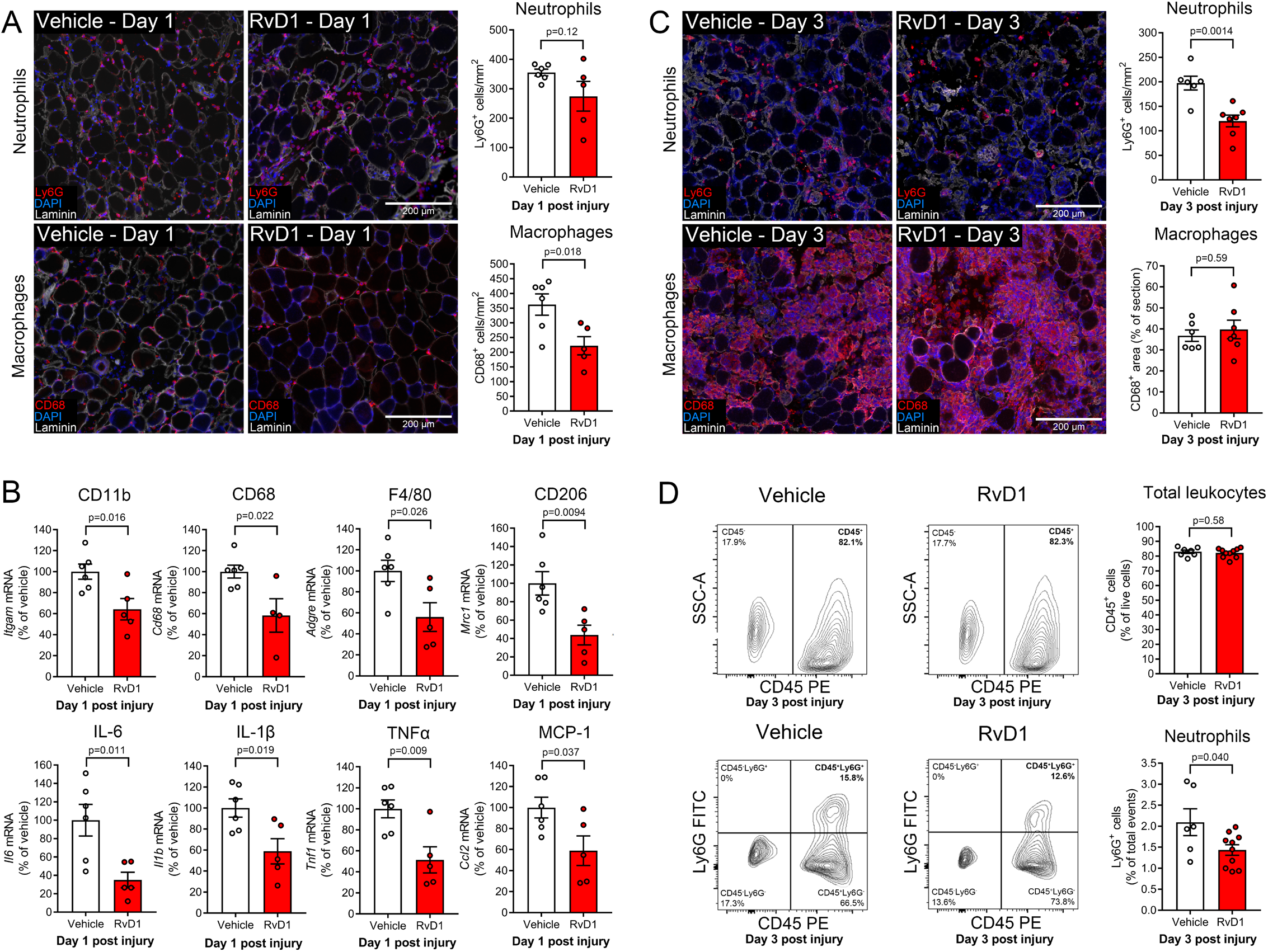
Resolvin D1 limits inflammation and expedites its resolution following muscle injury: A: Female C57BL/6 mice (4-6 mo) received bilateral intramuscular injection of the tibialis anterior (TA) muscle with 50 μL of 1.2% barium chloride (BaCl_2_) to induce myofiber injury. Mice were treated with a single dose of resolvin D1 (RvD1, 100 ng) or vehicle (0.1% ethanol) via intraperitoneal (IP) injection ~5 min prior to muscle injury. TA cross-sections were stained for polymorphonuclear cells (PMNs, Ly6G) or monocytes/macrophages (MΦ, CD68). Cell nuclei and the basal lamina were counterstained with DAPI and a laminin antibody respectively. Scale bars are 200 μm. B: Whole muscle mRNA expression of immune cell markers and inflammatory cytokines at 24 h post-injury as determined by RT-PCR. Gene of interest expression was normalized to beta-actin (*Actb*). C: Mice were treated for 72 h with daily IP injection of RvD1 (100 ng) or vehicle (0.1% ethanol) following muscle injury and TA cross-sections were stained for PMNs (Ly6G) or MΦ (CD68) at day 3 post-injury. Day 3 post-injury representative images show examples of regions of muscle not yet infiltrated by MΦ where lingering PMNs remained (top panels) and MΦ rich regions which were largely devoid of PMNs (bottom panels). The entire tissue section was assessed for quantitative analysis. D: Single-cells isolated from pooled left and right TA muscles at day 3 post-injury were analyzed by flow cytometry for CD45-PE and Ly6G-FITC. Dead cells were excluded from analysis using Live/dead fixable violet. Bars show the mean ± SEM of 5-10 mice per group with dots representing data from each individual mouse (biological replicates). P-values were determined two-tailed unpaired t-tests.

### Systemic Resolvin D1 Treatment Expedites Clearance of Intramuscular PMNs

To investigate the effect of RvD1 on resolution phase of the acute inflammatory response, we treated mice daily with IP injection of RvD1 for 72 hours during recovery from BaCl_2_ induced muscle injury. RvD1 treated mice showed ~40% fewer intramuscular PMNs (Ly6G^+^ cells), while the relative area of the muscle cross-section containing CD68^+^ staining was similar between groups, (~40% of tissue area) (Figure 3C). To confirm this apparent effect of RvD1 to specifically reduce muscle PMNs, TA muscles were collected at day 3 post-BaCl_2_ injury and the intramuscular single-cell population was isolated for analysis by flow cytometry (Figure 3D). Approximately 80% of the live single-cells within injured muscle stained positive for the pan leukocyte marker CD45 and RvD1 treatment did not impact total intramuscular leukocytes. Nevertheless, the relative proportion of gated live CD45^+^ cells that co-stained positive for the PMN marker Ly6G was lower in RvD1 treated mice (Figure 3D). This was accompanied by a parallel increase in the relative proportion of intramuscular CD45^+^Ly6G^−^ cells, the vast majority of which are most likely MΦ at this time-point.

### Resolvin D1 Directly Stimulates MΦ Phagocytosis

MΦ play an important role in the active resolution of inflammation via their phagocytic uptake and removal of pathogens, cellular debris, and apoptopic PMNs. Therefore, we assessed whether RvD1 could directly influence MΦ phagocytic activity *in-vitro* (Figure 4A). Treatment with RvD1 at doses between 10-100 nM stimulated phagocytosis of *E. Coli* Bio Particles by GM-CSF derived mouse bone marrow MΦ (BMMs) (Figure 4B). RvD1 (1-10 nM) also stimulated *E. coli* phagocytosis by human peripheral blood monocyte (PBMC) derived M1 polarized MΦ, although higher doses of RvD1 (100 nM) were apparently refractory in this cell type (Figure 4C).

**Figure 4.**
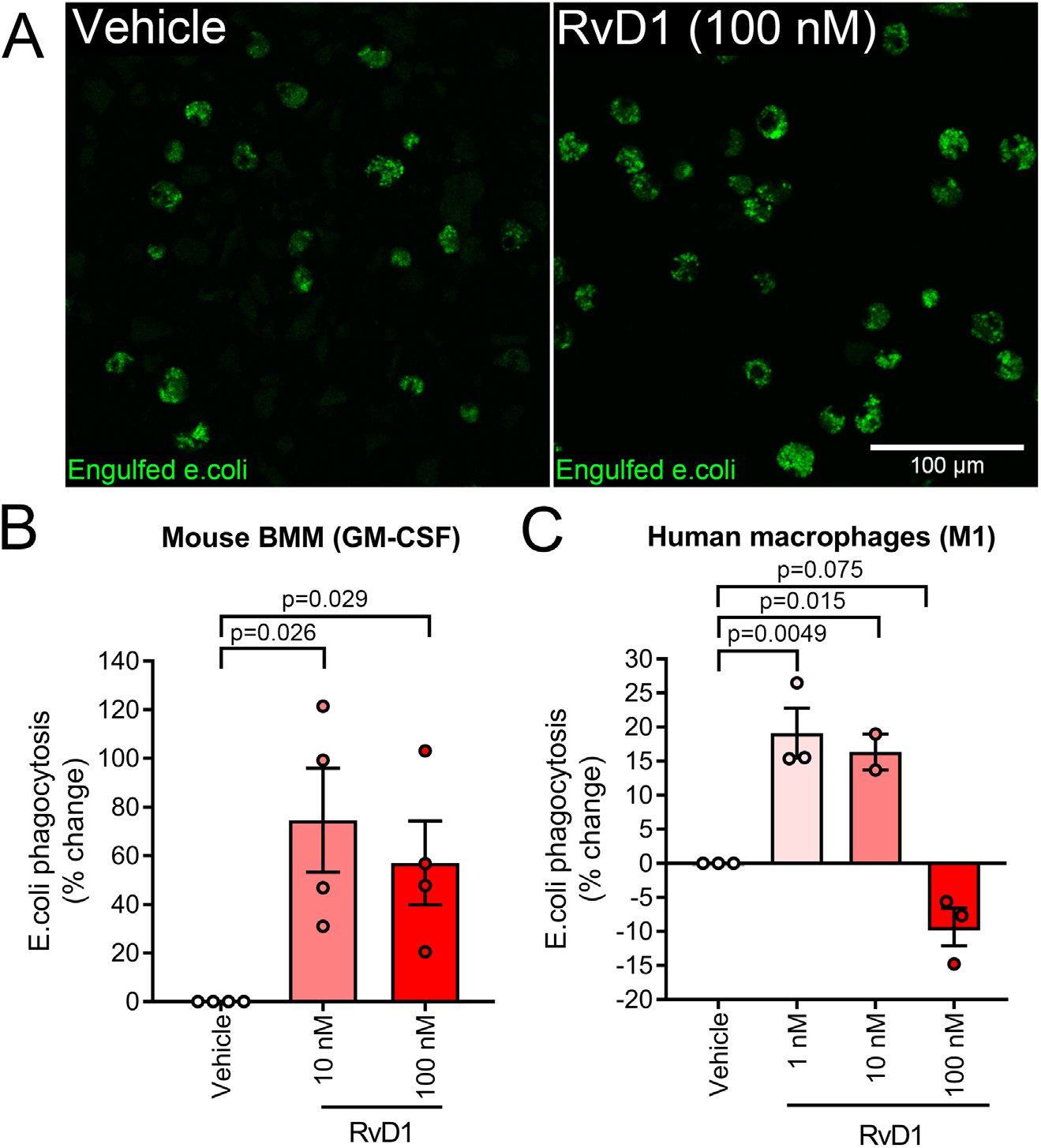
Resolvin D1 enhances macrophage phagocytosis: A: Bone marrow-derived macrophages (BMM) from female C57BL/6 mice (4-6 mo) were obtained by culturing myeloid precursor cells for 7 days in the presence of 20 ng/mL GM-CSF. GM-CSF derived BMMs were then pre-treated for 15 minutes with resolvin D1 (RvD1,100 nM) prior to incubation with pHrodo Green *E. Coli* Bio Particles for 60 min at 37°C in the continued presence of RvD1. B: Quantification of phagocytosis by mouse GM-CSF BMMs treated with RvD1 (10-100 nM) as determined using a 96-well fluorescent plate reader (excitation/emission of 509/533 nm). C: The effect of increasing doses of RvD1 (1-100 nM) on phagocytosis by human peripheral blood mononuclear cell (PBMC) derived M1 polarized macrophages. Data is presented at the percentage change in fluorescence intensity relative to matching vehicle treated macrophages from the same host. Bars show the mean ± SEM of cells originating from 3-4 different donors with dots with representing data from each host mouse or human subject (biological replicates). P-values were determined by one-way ANOVA followed by Holm-Sidak post-hoc tests with vehicle treated cells serving as a control group.

### Expedited Resolution of Muscle Inflammation Supports Regenerating Myofiber Growth

To test whether stimulating resolution influenced myofiber regeneration, mice were treated daily with systemic (IP) administration of RvD1 and TA muscles were collected at day 5 of recovery from BaCl_2_ injury. Hematoxylin and eosin (H&E) staining of TA muscle cross-sections did not reveal any gross histological differences in overall muscle morphology between vehicle and RvD1 treated mice (Figure 5A). Thus, RvD1 did not appear to obviously perturb normal muscle regeneration as has been reported previously with systemic NSAID treatment (20–25). To more precisely measure the extent of myofiber regeneration, tissue sections were stained with an antibody against embryonic myosin heavy chain (eMHC) (Figure 5A, Supplemental Figure 3). RvD1 did not influence the absolute or relative number of eMHC^+^ fibers within the regenerating TA, but did increase the average cross-sectional area (CSA) of the regenerating (eMHC^+^) myofiber population (Figure 5B).

**Figure 5.**
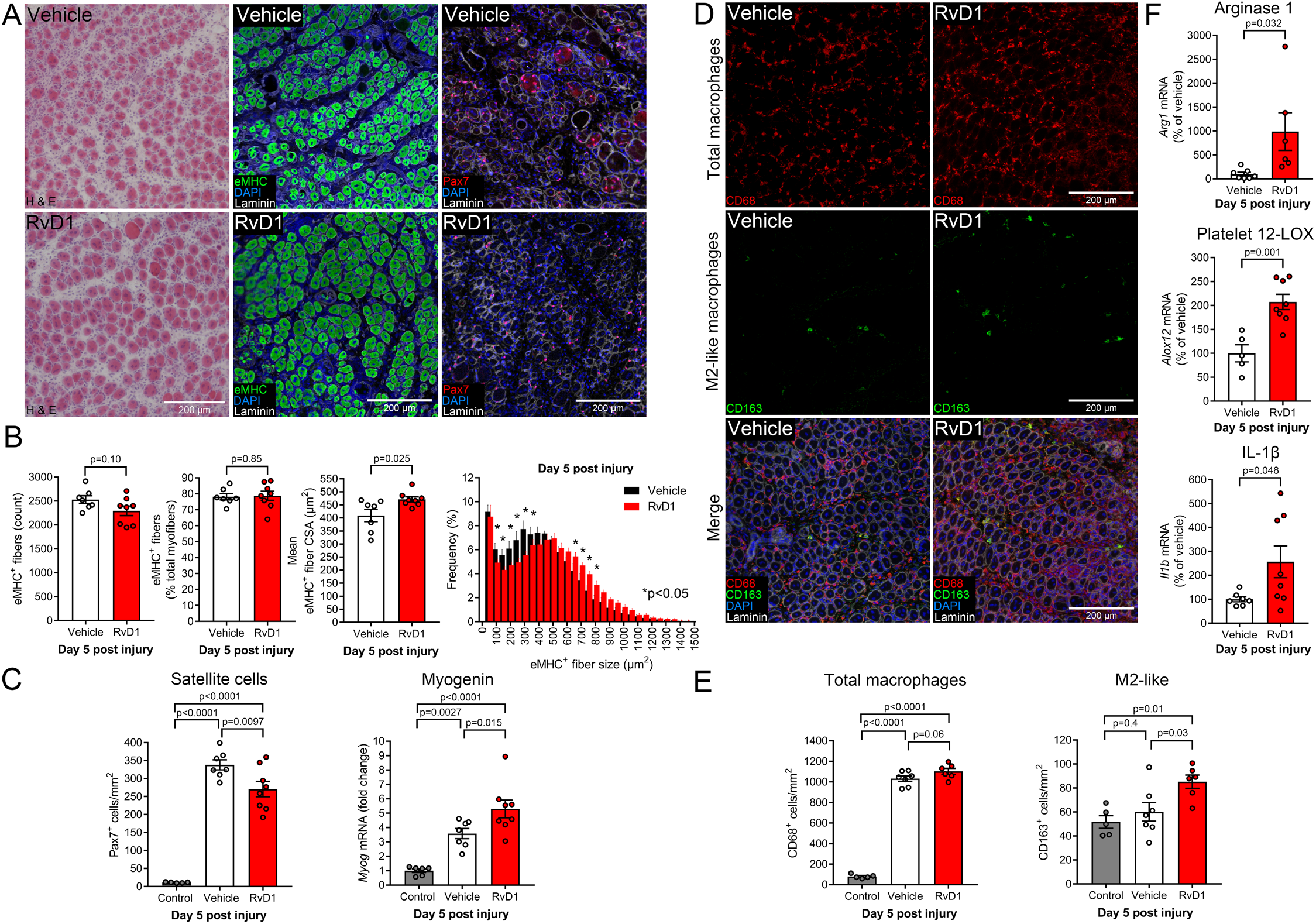
Resolvin D1 enhances myofiber regeneration by modulating muscle stem cell and macrophage responses to muscle injury: A: Female C57BL/6 mice (4-6 mo) received bilateral intramuscular injection of the tibialis anterior (TA) muscle with 50 μL of 1.2% barium chloride (BaCl_2_) to induce myofiber injury. Mice were then treated daily with resolvin D1 (RvD1, 100 ng) or vehicle (0.1% ethanol) by intraperitoneal (IP) injection for five days with the first dose administered ~5 min prior to injury. TA cross-sections were stained for hematoxylin & eosin (H&E), embryonic myosin heavy chain (eMHC), or Pax7 to stain muscle stem cells (satellite cells, MuSCs). Cell nuclei and the basal lamina were counterstained with DAPI and a laminin antibody respectively. Scale bars are 200 μm. B: Quantitative analysis of total regenerating (eMHC^+^) myofiber number, relative eMHC^+^ fiber number (as % of total fibers), mean eMHC^+^ fiber cross-sectional area (CSA), and the percentage frequency distribution of the CSA of the eMHC^+^ fiber population. C: Quantification of MuSC number (Pax7^+^/DAPI^+^ nuclei) and whole muscle mRNA expression of the myogenic regulatory factor myogenin at day 5 post-injury. D: Cross-sections of TA muscles at day 5 post-injury were stained for total macrophages (MΦ, CD68) and M2-like MΦ (CD163). E: Quantification of total intramuscular MΦ (CD68) and M2-like MΦ (CD163^+^ cells). F: Whole muscle mRNA expression of MΦ-related genes including arginase-1 (Arg1), platelet-type 12-lipoxygenase (12-LOX), and interleukin 1 beta (IL-1β) as determined by RT-PCR. Gene of interest expression was normalized to beta-actin (*Actb*). Bars show the mean ± SEM of 5-8 mice per group with dots representing data for each individual mouse (biological replicates). P-values were determined by one-way ANOVA followed by pair-wise Holm-Sidak post-hoc tests (panels C and E) or by two-tailed unpaired t-tests (panels B and F).

### Immunoresolvent Treatment Modulates Muscle Stem Cell Responses to Injury

Skeletal muscle regeneration is predominantly, if not entirely, dependent on the function of resident MuSCs (43). Therefore, we also stained muscle cross-sections with an antibody against the MuSC marker Pax7 (Figure 5A). The number of Pax7^+^ cells increased ~20 fold at day 5 following BaCl_2_ injury and RvD1 treatment reduced MuSC number (Figure 5C). Since myogenic regulatory factors control the activation, proliferation and differentiation of MuSCs, we questioned whether RvD1 influenced myogenic gene expression. Whole TA muscle mRNA expression of myogenin increased markedly at day 5 post-injury, and increased even further in mice that received RvD1 (Figure 5C). Taken together with the effect of RvD1 to increase regenerating myofiber size, a lower muscle Pax7^+^ cell density at day 5 post-injury may thus be interpreted as RvD1 stimulating myogenic commitment, differentiation, and/or fusion of MuSCs with regenerating myofibers.

### Resolvin D1 Shifts Intramuscular MΦ Activation State

Many MΦ persisted in muscle at day 5 post-BaCl_2_ injury and RvD1 did not influence the total number of muscle MΦ (Figure 5E), or muscle mRNA expression of MΦ markers including CD11b, CD68, F4/80 or CD206 (data not shown). Only a small proportion of intramuscular MΦ at day 5 post-injury co-expressed the CD163 antigen (≤5%) (Figure 5D). Nevertheless, RvD1 treated mice had greater numbers of CD163^+^ MΦ in regenerating muscle (Figure 5E). RvD1 also increased muscle mRNA expression of the M2 MΦ marker arginase-1 (Arg-1) and platelet-type 12-lipoxygenase (12-LOX), a key SPM biosynthetic enzyme that is highly expressed by mature MΦ (Figure 5F). Expression of the MΦ related pro-inflammatory cytokine IL-1β was also ~3 fold higher in regenerating muscle of RvD1 treated mice (Figure 5F), although other M1-related cytokines such as IL-6, MCP-1 and TNFα were not affected (data not shown).

### Improved Recovery of Muscle Function in Resolvin D1 Treated Mice

BaCl_2_ induced muscle injury resulted in a ~40% decrease in nerve-stimulated *in-situ* maximal isometric contractile force (P_o_) at day 14 post-injury and much of force deficit persisted when normalized to muscle size (specific force, sP_o_) (Figure 6A). Treatment with RvD1 improved recovery of P_o_ by ~15%, but did not significantly influence recovery of sP_o_. Histological analysis of regenerating muscles showed that RvD1 did not influence muscle fiber type composition, but the size of fast-twitch type IIb fibers was larger in RvD1 treated mice (Figure 6C). There was also an improved recovery of overall TA muscle size (CSA) in mice receiving RvD1 treatment (Figure 6D). ~60% of the muscle fibers contained centrally located myonuclei at this time-point. RvD1 did not influence the absolute or relative number of these regenerating myofibers but regenerating myofiber CSA was larger in mice treated with RvD1 (Figure 6D). Regenerating muscles still contained ~3-fold more MΦ than uninjured TAs (Figure 6E). Only a minority (~20%) of these CD68^+^ cells expressed CD163, however, suggesting that even by 14 days post-BaCl_2_ injury, the bulk of muscle MΦ are not M2-like in the traditional sense. RvD1 did not impact the number of intramuscular CD163^+^ MΦ, but did reduce total MΦ numbers back towards the numbers typically seen in uninured muscle (Figure 6G).

**Figure 6.**
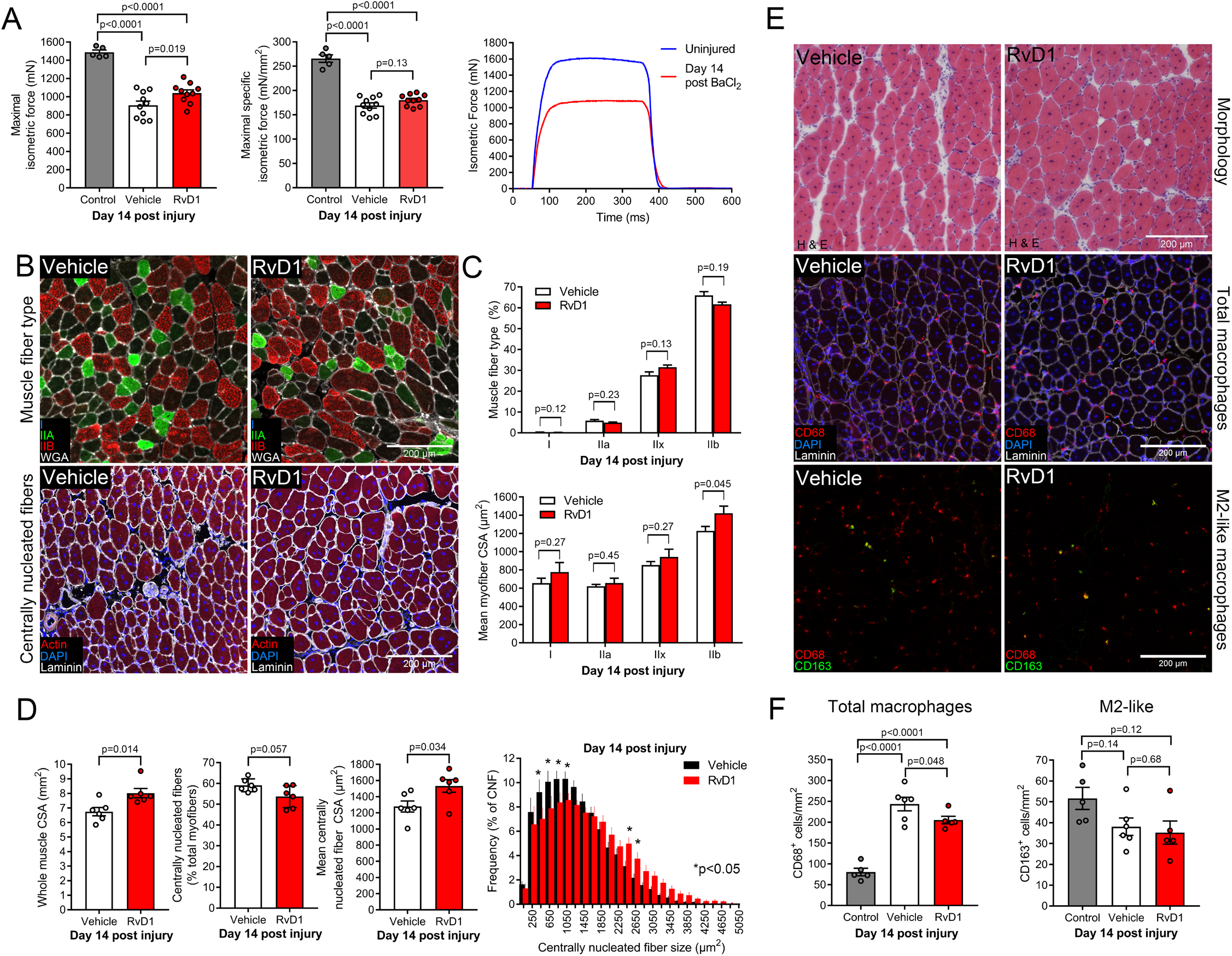
Resolvin D1 improves recovery of isometric muscle strength: A: Female C57BL/6 mice (4-6 mo) received bilateral intramuscular injection of the tibialis anterior (TA) with 50 μL of 1.2% barium chloride (BaCl_2_) to induce myofiber injury. Mice were then treated daily with intraperitoneal (IP) injection of resolvin D1 (RvD1, 100 ng) or vehicle (0.1% ethanol) for 14 days with the first dose administered ~5 min prior to injury. TA muscle function was then tested for maximal isometric nerve-stimulated *in-situ* contractile force (P_o_) with age and gender matched mice serving as uninjured controls. Absolute maximal isometric force (mN) generated by the TA muscle was measured and used to calculate maximal specific isometric contractile force (sP_o_, mN/mm^2^). Representative force traces obtained from uninjured and injured TA muscles are shown. B: TA cross-sections were stained with conjugated phalloidin to label the total muscle fiber population (actin filaments) or for muscle fiber type with antibodies against type myosin heavy chain I, IIa, and IIb. Type IIx fibers remain unstained (black) and are identified by lack of fluorescent staining. Cell nuclei and the basal lamina were counterstained with DAPI and a laminin antibody respectively. Scale bars are 200 μm. C: Quantitative analysis of percent muscle fiber type composition and mean fiber cross-sectional area (CSA) split by muscle fiber type as determined by MuscleJ software. D: Quantification of overall TA muscle CSA, the percentage of centrally nucleated (regenerating) myofibers, mean regenerating myofiber CSA, and percent frequency distribution of regenerating fiber CSA as determined by MuscleJ software. E: TA cross-sections were stained for hematoxylin & eosin (H&E), total macrophages (MΦ, CD68^+^ cells), and M2-like MΦ (CD163^+^ cells). Scale bars are 200 μm. F: Quantification of the histological presence of total muscle MΦ and M2-like MΦ. Cell counts were performed manually throughout the entire muscle cross-section and then normalized to tissue surface area as determined by MuscleJ software. Bars show the mean ± SEM of 5-10 mice per group with dots representing data from each individual mouse (biological replicates). P-values were determined by one-way ANOVA followed by pair-wise Holm-Sidak post-hoc tests (panels A and F) or two-tailed unpaired t-tests (panels C and D).

### Resolvin D1 Minimally Impacts the Global MuSC Transcriptome, but May Specifically Modulates Genes Related to Muscle-Immune Interactions

In order to gain mechanistic insight into the underlying mechanisms by which RvD1 enhanced muscle regeneration, we performed transcriptome wide profiling of gene expression of MuSCs isolated from day 3 post-BaCl_2_ injured TA muscles via fluorescence activated cell sorting (FACS) (CD45^−^, CD11b^−^, CD31^−^, Sca-1^−^, Ter-119^−^, CXCR4^+^, Integrin beta-1^+^ cells) followed by RNA-sequencing. The absolute MuSC yield from injured TA muscles was lower in RvD1 treated mice (Figure 7A), while TA muscle mass was not affected (Figure 7B), indicating relative muscle MuSC number was reduced in RvD1 treated mice at this time-point (Figure 7C). Overall the global transcriptomic profile of isolated MuSCs from vehicle and RvD1 treated mice at day 3 post-injury was very similar (Figure 7D), and differed markedly from uninjured MuSCs (Supplemental Figure 4A). Nevertheless, assessment of differentially expressed genes (>2-fold change, irrespective of p-value) between RvD1 and vehicle treated mice did reveal an enrichment of genes associated with response to wounding (76 genes), the inflammatory response (54 genes), response to external stimulus (98 genes), and cytokine secretion (17 genes), amongst others (Figure 7E). A heat map of the 100 most different annotated genes in isolated MuSCs between vehicle and RvD1 treated mice is shown in Supplemental Figure 4B.

**Figure 7.**
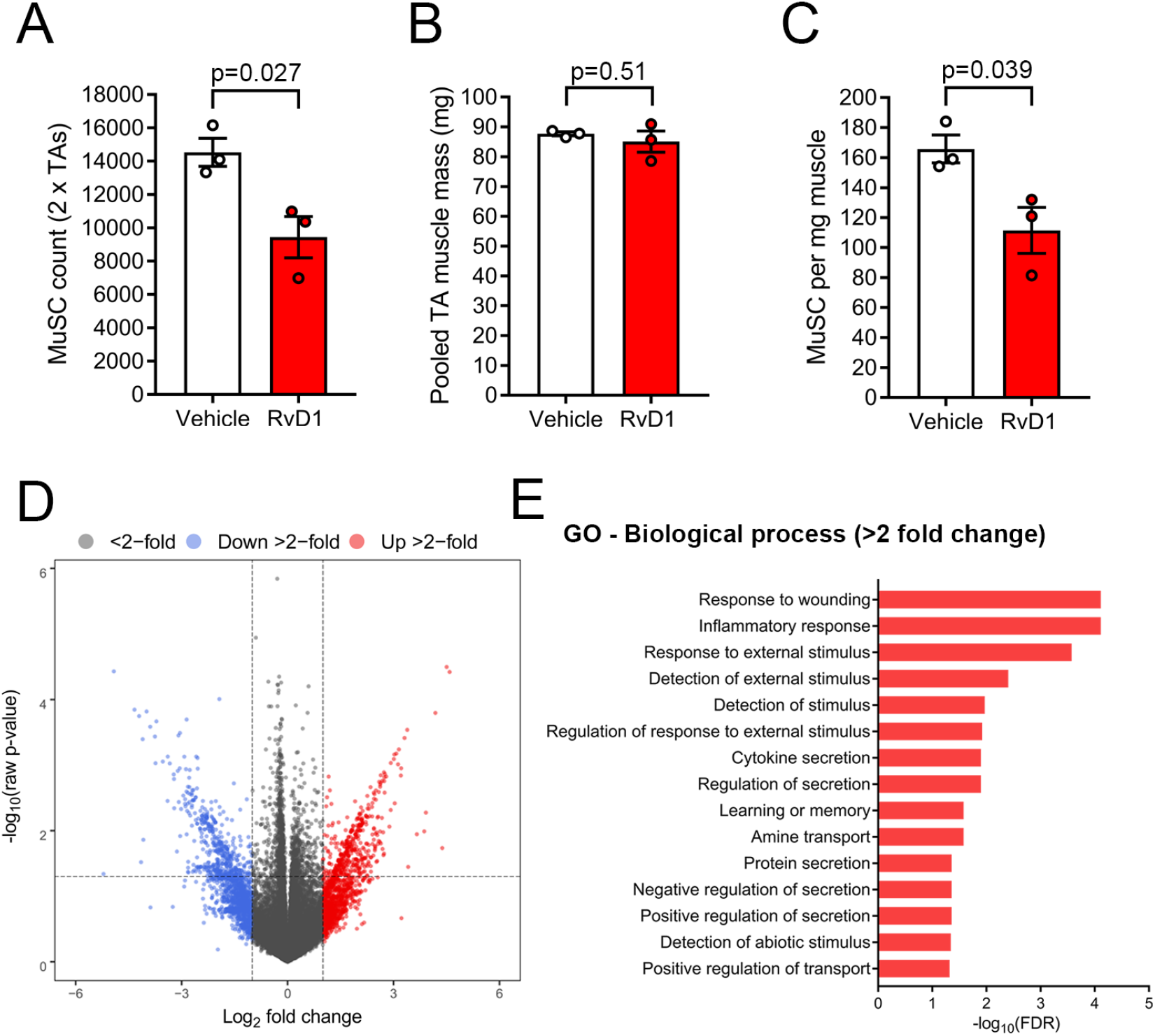
Resolvin D1 minimally impacts the global transcriptomic response of muscle stem cells to injury, but may specifically modulate genes related to muscle-immune cell interactions: Female C57BL/6 mice (4-6 mo) received bilateral intramuscular injection of the tibialis anterior (TA) muscle with 50 μL of 1.2% barium chloride (BaCl_2_) induce myofiber injury. Mice were treated with daily intraperitoneal (IP) injection with resolvin D1 (RvD1, 100 ng) or vehicle control (0.1% ethanol) for 72 h with the first dose administered ~5 min prior to injury. Both TA muscles were collected at day 3 post-injury and pooled. Muscle stem cells (satellite cells, MuSCs) were isolated by fluorescence-activated cell sorting (FACS), and transcriptome wide profiling of the isolated MuSC population was performed by RNA-sequencing. A: Total MuSC yield from day 3 post-injury TA muscles. B: Pooled mass of TA muscles used for MuSC isolation. C: Relative MuSC yield. D: Volcano plot of overall RNA-seq data with each dot representing a single gene, positive Log_2_ fold changes (FC) indicating induction and negative Log_2_ FC indicating suppression in response to RvD1 treatment. Genes induced >2 FC (+1 Log_2_ FC) are colored red and those suppressed >2 FC (−1 Log_2_ FC) are colored blue. E: Gene ontology (GO) enrichment and respective false discovery rates (FDR) for genes up or downregulated >2 FC (irrespective of p-value) in response to RvD1. Bars show the mean ± SEM of 3 mice per group with each dot representing data from an individual mouse (biological replicates). P-values were determined two-tailed unpaired t-tests.

### Physiological Doses of Resolvin D1 Neither Promote nor Perturb *In-Vitro* Myogenesis

We also tested whether RvD1 could directly modulate muscle cell growth *in-vitro*. Treatment with a single dose of 100 nM RvD1 [based on established bioactivity in murine MΦ (Figure 4)] at the time of myogenic differentiation did not influence cellular density, percentage of myoblast differentiation, or fused myotube size (Figure 8A). Replenishing the culture media with fresh RvD1 every 24 hours, in an attempt to account for any potential instability of RvD1 also did not influence *in-vitro* myogenesis (data not shown). Since NSAIDs have direct suppressive effects on *in-vitro* myogenesis (61), we questioned whether higher doses of RvD1 would also interfere with myogenesis. The non-specific COX pathway inhibitors ibuprofen and indomethacin, as well as the COX-2 selective inhibitor NS-398, impaired myotube development in a dose-dependent manner (Supplemental Figure 5). Maximally effective doses of NS-398 (50 μM), ibuprofen (500 μM), and indomethacin (200 μM) all quantitatively reduced indices of myoblast density, percentage differentiation, and fused myotube diameter (Figure 8B). In contrast, muscle cells treated with RvD1 at doses of 0.1-1 μM, showed no such deleterious effects and a modest yet statistically significant increase in fused myotube size (+5%) was observed with 1 μM RvD1 (Figure 8B).

**Figure 8.**
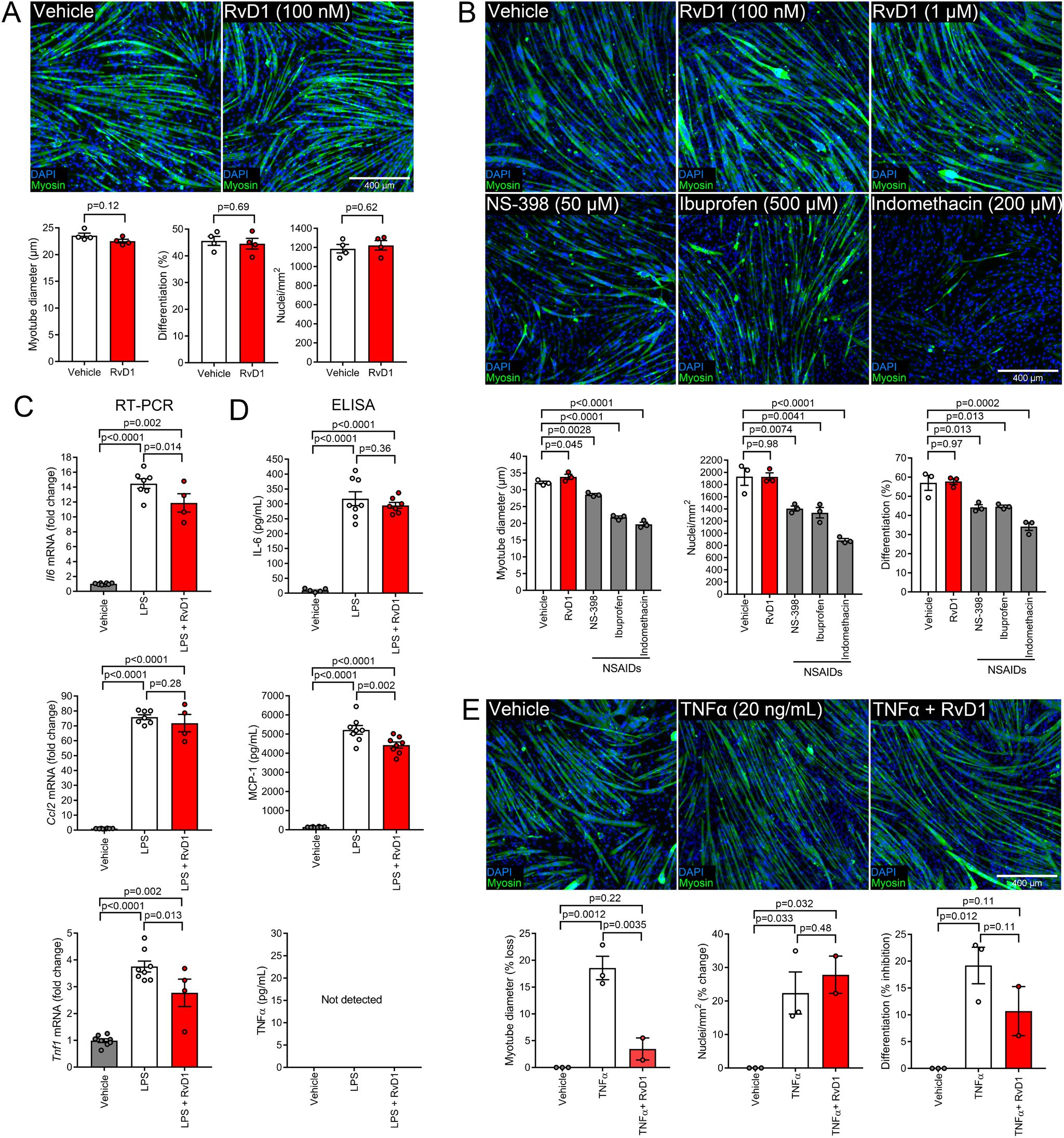
Resolvin D1 neither enhances nor perturbs basal *in-vitro* myogenesis, but directly modulates myokine production and protects muscle cells against the deleterious effects chronic inflammation: A: Murine C2C12 myoblasts were treated with resolvin D1 (RvD1, 100 nM) at onset of myogenic differentiation. At 3-days post-differentiation, myotubes were fixed in 4% paraformaldehyde and stained for sarcomeric myosin. Cell nuclei were counterstained with DAPI. Quantitative analysis was performed on 6 non-consecutive fields of view per well to determine mean myotube diameter, percentage of myogenic differentiation, and overall cell density. B: Confluent C2C12 myoblasts were induced to differentiate in the presence of RvD1 (0.1-1 μM), or non-steroidal anti-inflammatory drugs (NSAIDs) including NS-398 (50 μM), ibuprofen (500 μM), and indomethacin (200 μM). C: C2C12 myotubes at day 3 post-differentiation were pre-treated with RvD1 (100 nM) for 30 min before stimulation with lipopolysaccharide (LPS, 100 ng/mL) for 3 h in the continued presence of RvD1. Cellular mRNA expression of cytokines including interleukin-6 (IL-6), monocyte chemoattractant protein 1 (MCP-1), and tumor necrosis factor alpha (TNFα) was determined by RT-PCR. D: C2C12 myotubes at day 3 post-differentiation were pre-treated with RvD1 (100 nM) for 30 min and then stimulated with LPS (100 ng/mL) for 24 h in the continued presence of RvD1. Conditioned culture media was collected from the cells and analyzed for concentrations of the cytokines IL-6, MCP-1 and TNFα by ELISA. E: Confluent C2C12 myoblasts were induced to differentiate in the presence of exogenous TNFα (20 ng/mL), with or without RvD1 co-treatment (100 nM). At day 3 post-differentiation, myotubes were fixed, stained, and quantified as described in panel A. Scale bars are 400 μm. Bars show the mean ± SEM of 3-8 replicates per group with each dot representing data for a single independent culture well which was considered as a biological replicate. P-values were determined by two-tailed unpaired t-tests (panel A) or one-way ANOVA followed by pair-wise Holm-Sidak post-hoc tests (panels B-E).

### Resolvin D1 Directly Limits Myokine Production and Protects Muscle Cells Under Conditions of Chronic Inflammatory Stress

RvD1 is well established to suppress cytokine production by a variety of cell types, most notably MΦ (62–67). Therefore, we questioned whether RvD1 could directly modulate inflammatory cytokines in muscle cells (myokines). Exposure of C2C12 myotubes to lipopolysaccharide (LPS) markedly increased production of IL-6, MCP-1 and TNFα at the gene and protein level (Figure 8C-D). Pre-treatment with RvD1 blunted LPS stimulated (3 h) mRNA expression of IL-6 and TNFα, but did not influence MCP-1 (Figure 8C). Following more prolonged treatment (24 h), RvD1 also reduced LPS stimulated secretion of MCP-1 (Figure 8D). Despite this, prolonged RvD1 treatment did not statistically reduce IL-6 secretion and TNFα was not present at detectable concentrations from LPS stimulated C2C12 cells with or without RvD1 treatment. In order to determine whether RvD1 may directly influence myogenesis under conditions of chronic non-resolving inflammation, C2C12 myoblasts were also induced to differentiate in the presence of the pro-inflammatory cytokine TNFα (20 ng/mL) (Figure 8E). Long-term TNFα exposure reduced the number of myoblasts successfully undergoing myogenic differentiation and lead to formation of thin elongated myotubes. Co-treatment with RvD1 (100 nM) protected against the deleterious effect of TNFα on developing myotube size, but did not rescue TNFα induced effects on myoblast cellular density or myogenic differentiation (Figure 8E).

## Discussion

Here we investigated the role of specialized pro-resolving mediators (SPMs) in inflammatory and adaptive responses to muscle damage and tested the efficacy of systemic immunoresolvent treatment as a novel therapeutic strategy to the treatment of muscular injuries. SPMs were locally produced in muscle in response to both chemical injury and functional overload, coinciding with peak MΦ infiltration, clearance of tissue PMNs, and the onset of adaptive myofiber remodeling. Daily systemic treatment with the native SPM RvD1 as an immunoresolvent suppressed muscle inflammatory cytokines, expedited tissue PMN clearance, and shifted intramuscular MΦ to a more M2-like activation state. These immunomodulatory effects were associated with enhanced regenerating myofiber growth and improved recovery of muscle strength. While RvD1 had little direct impact on myogenesis *in-vitro*, it directly suppressed myokine production and enhanced MΦ phagocytic activity, indicative of both muscle direct and indirect actions. These findings show that SPMs play an important supportive role in myofiber growth and regeneration and highlight their therapeutic potential in the context of muscle injuries as a novel alternative to classic anti-inflammatory interventions (e.g. NSAIDs), which are known to inhibit endogenous cellular regenerative mechanisms.

Intramuscular injection of BaCl_2_ results in wide-spread myofiber necrosis and a robust inflammatory response (68–70). In this model, restoration of muscle function is dependent on *de-novo* muscle fiber formation via the proliferation, differentiation, and fusion of MuSCs (myofiber regeneration) (43). In contrast, functional muscle overload induced by synergist ablation results in mild myofiber damage, and ensuing adaptive tissue remodeling occurs mainly as a result of compensatory growth of pre-existing muscle cells (myofiber hypertrophy) (71). We show here that both interventions resulted in rapid infiltration of muscle by PMNs, their subsequent disappearance, and a persistent intramuscular MΦ presence. In both models, there was an initial increase in expression of inducible COX-2, leading to increased intramuscular concentrations of classical pro-inflammatory eicosanoids. COX pathway products including PGE_2_ (72, 73), PGF_2α_ (74–76) and PGI_2_ (77, 78) have all been previously implicated in stimulating various stages of skeletal muscle regeneration. PGE_2_ also plays a key role in signaling the onset of the resolution phase by inducing transcription of 15-LOX, a phenomenon termed lipid mediator class switching (79). Consistent with this concept, expression of 5-, 12- and 15-LOX, as well as intramuscular concentrations of many LOX derived monohydroxylated fatty acid intermediates in SPM biosynthesis pathways increased locally during muscle regeneration. COX-2 exhibited a bimodal response, which is consistent with the proposed dual role of prostaglandins in both the induction and resolution of acute inflammation (80). Heightened local 5- and 15-LOX expression and increased tissue abundance of many SPM pathway markers were also observed following functional overload of the rat plantaris muscle for the first time.

Bioactive SPMs themselves were generally not present at detectable concentrations in uninjured muscle, but intramuscular protectin D1 and maresin 1 were detected in response to both BaCl_2_ injury and functional overload, while resolvin D6 and lipoxin A_4_ were also detected in the latter model. SPMs and/or their monohydroxylated pathway markers have also recently been found to increase in mouse muscle following experimental ischemia (81), intramuscular cardiotoxin injection (38), and eccentric exercise (38), as well as in human muscle biopsies following an acute bout of damaging muscular contractions (39). Furthermore, acute exercise transiently increases SPMs in human blood (41, 42), and repeated exercise training was recently found to prime murine MΦ to release greater amounts of RvD1 in response to a subsequent inflammatory challenge (82). PMNs (Ly6G^+^), inflammatory (Ly6C^hi^F4/80^lo^) monocytes, and reparative (Ly6C^lo^F4/80^hi^) MΦ populations isolated from cardiotoxin injured mouse muscle were also recently found to display distinct time-dependent modulation of bioactive lipid mediators (38). Overall, these findings show that dynamic regulation of the SPM bioactive metabolome is an endogenous mechanism that is conserved across multiple different species and models of muscle damage.

We did not detect endogenous RvD1 within muscle before or after injury, but we did observe increased expression of the enzymatic machinery required in RvD1 biosynthesis (15- and 5-LOX) (48, 49), as well as intramuscular concentrations of 17-HDoHE, the primary 15-LOX intermediate product of n-3 DHA produced during D-series resolvin biosynthesis (48, 49). We (41) and others (42) previously found that circulating RvD1 does increase following strenuous exercise. Notably, primary monohydroxylated 5-, 12, and 15-LOX metabolites were far more abundant in muscle homogenates in the current study than bioactive SPMs themselves. This finding is consistent with our prior study of metabolipidomic profiling of human muscle biopsies (39), as well as recent reports by others in mouse muscle (38, 81). This suggests that the lipid metabolome of muscle may predominantly reflect intracellular metabolites whereas bioactive SPMs may be relatively enriched in the extracellular environment where they bind to their cell surface receptors. Another possibility is that bioactive SPMs in tissues are rapidly converted to downstream enzymatic inactivation products. Consistent with this concept, we observed 8-oxo-RvD1 [a bioactive 15-hydroxy prostaglandin dehydrogenase metabolite of RvD1 (48)] in rat muscle 3 days post-synergist ablation at levels ~2-fold higher than baseline.

Prior studies have shown that PMNs can directly damage muscle cells *in-vitro* (83), and that blockade of PMN influx following muscle damage, either by knockout of CD18 (7) or depletion of circulating PMNs (84), protects muscle from leukocyte-induced secondary injury. Therefore, limiting PMN recruitment and hastening their removal has been proposed to be organ protective in the context of sterile skeletal muscle injury. A class defining action of SPMs is the selective inhibition of further PMN recruitment to the site of inflammation (12, 13). Indeed, systemic injection of resolvins displays similar potency in reducing PMN infiltration in the murine peritonitis model as the NSAID indomethacin (13, 48, 49). On this basis, we hypothesized that treatment with RvD1 at the time of BaCl_2_ injury would limit the initial appearance of PMNs within injured muscle. RvD1 treatment did not influence peak intramuscular PMN, but did reduce intramuscular PMNs at day 3 of recovery. In prior studies, RvD1 also did not reduce peak PMN infiltration of cardiac muscle following myocardial infarction, and rather specifically accelerated PMN egress from the infarcted left ventricle (34, 35). Overall, these data are consistent with the notion that in addition to their anti-inflammatory actions, SPMs actively promote PMN clearance, thus accelerating resolution back to a non-inflamed state (85). One potential mechanism, is the ability of SPMs to stimulate MΦ mediated phagocytic uptake and removal of apoptopic PMNs (86). Consistent with prior studies (87), RvD1 proved to be a potent stimulator of MΦ phagocytic activity in our hands. Indeed, administration of RvD2 at the peak of inflammation (24 h) following ligation of the femoral artery reduced PMN numbers in the hamstring musculature of mice when assessed 48 h later (day 3 post-ischemia) (81). Notably, traditional anti-inflammatory drugs such as the NSAIDs, which lack the pro-resolving bioactivity that we and others have observed for SPMs, can inadvertently delay timely resolution of inflammation by limiting monocyte recruitment and interfering with MΦ mediated clearance of tissue PMNs (80).

One key molecular mechanism by which resolvins function is by counteracting production of pro-inflammatory cytokines (13, 49). Indeed, we found that systemic treatment with RvD1 at the time of muscle injury suppressed local cytokine expression. RvD1 also directly reduced LPS stimulated mRNA expression of IL-6 and TNFα, as well as secretion of MCP-1 protein by C2C12 myotubes cultured *in-vitro*. Consistent with these observations, n-3 EPA derived resolvin E1 (RvE1) directly inhibits cytokine production by muscle cells (88). While the overall impact of systemic RvD1 treatment on the global MuSC transcriptome of following injury was minimal in the current study, clusters of genes with strong relevance to muscle-immune interactions such as those involved in the response to wounding, the inflammatory response, and cytokine secretion were amongst those most influenced by RvD1 based on magnitude of change. Collectively, these data suggest that resolvins may have some direct modulatory effects on the production and/or release of inflammatory mediators by cells resident to the musculature such as post-mitotic myofibers and resident MuSCs.

The chemokine MCP-1 is indispensable for recruitment of blood monocytes to the injured musculature (89). Given the suppressive effect of RvD1 on muscle MCP-1 expression, the initial reduction observed in the current study in intramuscular MΦ following RvD1 treatment is not surprising. Despite this, the recovery of muscle MΦ numbers to normal levels during the resolution phase suggests that RvD1 treatment may have either stimulated later monocyte recruitment or promoted expansion of tissue MΦ. Mice receiving RvD1 also displayed heightened muscle expression of the alternative MΦ activation marker arginase-1, as well as a greater number of intramuscular MΦ expressing the M2-like marker CD163. Overall, these data are consistent with prior studies showing that RvD1 can induce a M2-like polarization state of MΦ *in-vitro* (52, 67, 90). In turn, M2-polarized MΦ produce more SPMs than M1 activated MΦ (91–94), which may explain why muscle expression of platelet-type 12-LOX, a key SPM biosynthetic enzyme expressed by MΦ, was increased in muscle in response to RvD1 treatment in the current study. Whilst most M1 signature cytokines did not differ between vehicle and RvD1 treated mice during the regenerative phase, IL-1β, a classical pro-inflammatory cytokine, was ~3 fold higher in muscle of RvD1 treated mice at day 5 post-injury. This may potentially be explained by recent studies showing that resolution phase MΦ are neither classically nor alternatively activated, but possess certain aspects of both phenotypes (52, 95).

NSAIDs have been extensively shown to have deleterious effects on muscle repair (20–25). Given that SPMs also possess some anti-inflammatory actions, it was important to determine whether repeated daily immunoresolvent treatment would impact the efficiency of muscle regeneration. Overall, we found no evidence of a deleterious effect of RvD1 on any indices of myofiber regeneration *in-vivo*. Moreover, the well-established direct suppressive effects of NSAIDs on *in-vitro* myogenesis (21, 61) were clearly not shared by RvD1, even at high doses. Rather, RvD1 enhanced muscle repair in mice *in-vivo* as evidenced by increased regenerating muscle fiber size and improved recovery of muscle strength. Additionally, high doses of RvD1 directly promoted hypertrophy of developing myotubes *in-vitro* and lower doses could protect muscle cells from the deleterious effects of chronic exposure to pro-inflammatory stimuli, as previously shown with RvE1 (88). The contrasting effects of NSAIDs and RvD1 on muscle regenerative capacity may be explained by their varying mechanisms of action. Whereas NSAIDs decrease intramuscular MΦ infiltration (20), impair M2-polarization (22), and suppress MΦ phagocytosis (86), RvD1 had overall stimulatory effects on such MΦ responses. MΦ in general, and M2-like MΦ in particular, are thought to support muscle regeneration (96). Therefore, the beneficial effect of RvD1 on muscle regeneration may be attributable to direct positive interactions between intramuscular MΦ and resident myofibers and/or MuSCs. Consistent with this hypothesis, RvD1 modulated the MuSC response during muscle regeneration based on an accelerated fusion of proliferated Pax7^+^ cells and increased expression of the myogenic regulator factor myogenin. Phagocytosis of muscle cell debris is a key stimulatory cue triggering a MΦ polarization (4), and blocking phagocytosis has deleterious effects on both intramuscular MΦ phenotype transitions and myofiber regeneration (6). Therefore, the stimulatory effects of RvD1 on phagocytosis, intramuscular MΦ activity, and myofiber regeneration are likely to be complex and intrinsically linked.

Recently, a single intramuscular injection of resolvin D2 (RvD2) was reported to improve recovery of muscle mass and strength following muscle injury in mice (38). However, that study focused primarily on immunological outcomes and the underlying cellular and molecular basis for these apparent effects on muscle regeneration were not investigated. Moreover, the mice were depleted of resident MuSCs by irradiation, which is known to markedly deregulate the normal inflammatory and regenerative responses to muscle injury (97), as further evidenced by the marked weakness at day 14 of recovery from injury compared with that measured for the healthy young mice in the present study (900 vs. 200 mN P_o_) (38). Under the physiological relevant circumstances in which MuSCs play a well-established and indispensable role in *de-novo* myofiber formation (43), we observed that the beneficial effects of RvD1 on recovery of muscle size and strength were mechanistically attributable to hypertrophy of regenerating myofibers. This was accompanied by enhanced myogenic gene expression and an accelerated fusion of proliferated MuSCs, indicative of modulatory effects on this key resident myogenic stem cell population. Furthermore, transcriptome-wide profiling of MuSCs isolated from injured muscle showed that RvD1 modulated expression of clusters of genes relevant to muscle-immune interactions *in-vivo* and directly modulated myokine production *in-vitro*. In addition to recently reported effects of RvD2 on MΦ (38), we further show an impact of immunoresolvent treatment on intramuscular PMN clearance from injured muscle, which provides one more mechanism that may explain the therapeutic benefit of immunoresolvents in the context of muscle regeneration (7). In support of our findings, the pro-resolving peptide Annexin A1, a distinct agonist of the RvD1 receptor (FPR2/ALX), was also recently shown to be required for effective myofiber regeneration in healthy young mice (44).

The demonstrated benefit of systemic immunoresolvent treatment on muscle inflammation and regeneration in the current study has important implications for future therapeutic translation to human studies, especially given that RvD1 is orally bioavailable in mice (98). Moreover, the ability of daily dosing with RvD1 throughout the entire time-course of recovery from injury to ultimately improve muscle regeneration rather than compromise it, as is the case with daily systemic NSAID treatment (20–25), is in itself novel. This clear advantage of RvD1 treatment compared with NSAIDs is of great clinical importance given that such frequent repeated dosing would likely be required to effectively exploit the analgesic potential of SPMs for pain management (99, 100), which is a key therapeutic goal in the clinical treatment of soft tissue injuries. An additional novel finding of the current study is the demonstration for the first time that SPM biosynthetic circuits are also locally induced in response to overload of skeletal muscle, in close association with intramuscular immune cell transitions and the onset of the adaptive hypertrophy of pre-existing myofibers. In addition to their key supportive role in muscle regeneration, MΦ also regulate overload-induced myofiber hypertrophy (101) and myofiber re-growth during recovery from disuse (102). Thus, our observation of SPM biosynthesis in muscle with increased load suggests that immunoresolvents may also have potential therapeutic applicability to modulate adaptive responses to functional unloading and loading of muscle in physiologically and clinically relevant settings. Therefore, further studies should address the effects of immunoresolvents on muscle in models of adaptive remodeling of pre-existing post-mitotic myofibers such as limb immobilization or bed rest, and subsequent return to ambulation and rehabilitation. Immunoresolvents may also be an effective novel therapeutic in physiological and clinically relevant settings characterized by sustained non-resolving acute inflammatory responses and limited regenerative efficiency such as volumetric muscle loss (103). Finally, states of chronic unresolved inflammation which negatively impact upon muscle mass and myofiber regenerative capacity including aging (104), muscular dystrophies (19), and metabolic disease (e.g. obesity/diabetes) (105, 106) may also benefit from this novel therapeutic strategy and should be investigated.

In conclusion, SPMs are locally produced in response to myofiber damage and play an important role in the control of muscle inflammation, its active resolution, and cellular transitions of MΦ, MuSCs and myofibers to enable effective muscle tissue repair. These data implicate SPMs in controlling adaptive muscle remodeling and highlight the potential efficacy of systemic immunoresolving therapies as a novel treatment of muscular injuries which can enhance tissue regenerative capacity by stimulating endogenous resolution mechanisms.

## Materials and Methods

### Animal Handling

C57BL/6 mice and Sprague-Dawley rats were obtained from Charles River Laboratories and housed under specific pathogen free conditions with ad-libitum access to food and water. 4-6 month-old female mice were used for chemical muscle injury and 6-month-old male rats were used for functional muscle overload experiments.

### Chemical Induced Muscle Injury

Mice were anesthetized with 2% isoflurane and received bilateral intramuscular injection of the tibialis anterior (TA) muscle with 50 μL per limb of 1.2% BaCl_2_ in sterile saline (68–70). In some experiments mice randomized to receive sham injury via intramuscular injection of sterile saline alone instead of BaCl_2_. Mice were returned to their home cage to recover and monitored until ambulatory.

### Functional Muscle Overload

Synergist ablation was used to assess the inflammatory response to functional overload of the plantaris muscle (71). To minimize the impact of surgical manipulation on tissue inflammation, we used a mild protocol in which only the soleus/gastrocnemius tendon is surgically ablated. Rats were anesthetized with 2% isoflurane and preemptive analgesia provided by subcutaneous injection of buprenorphine (0.03 mg/kg) and carprofen (5 mg/kg). The skin overlying the posterior hind-limb was shaved and scrubbed with chlorhexidine and ethyl alcohol. A midline incision was made to visualize the gastrocnemius/soleus tendon and a full thickness tenectomy performed while leaving the plantaris tendon intact. The incision was closed using 4-0 Vicryl sutures. The procedure was then repeated on the contralateral limb to induce bilateral functional overload of both plantaris muscles. Following surgery, rats were returned to their cage to recover and monitored until ambulatory with free access to food and water. Postoperative analgesia was provided via subcutaneous injection of buprenorphine (0.03 mg/kg) at 12 h post-surgery and animals were monitored daily for any signs of pain or distress for 7 days. Age and gender matched rats served as non-surgical controls.

### Immunoresolvent Treatment

Resolvin D1 (RvD1, 7S,8R,17S-trihydroxy-4Z,9E,11E,13Z,15E,19Z-docosahexaenoic acid) was purchased from Cayman Chemicals (#10012554). Single use aliquots of RvD1 were prepared in amber glass vials (ThermoFisher, C4010-88AW) which were purged with nitrogen gas and stored at −80°C. On the day of use, the ethanol was evaporated to dryness under a gentle stream of nitrogen gas and RvD1 was re-suspended in sterile saline containing 0.1% ethanol. RvD1 stocks were handled in a darkroom and aqueous solutions of RvD1 were protected from light and used within 30 minutes of preparation. Mice were randomized to receive daily 100 μL intraperitoneal (IP) injections of either 100 ng of RvD1 or vehicle control (0.1% ethanol) with the first dose administered ~5 min prior to muscle injury. Mice were allowed to recover for up to two-weeks post-injury with daily IP injection of 100 ng of RvD1 or vehicle.

### Muscle Tissue Collection

Animals were euthanized via induction of bilateral pneumothorax while under deep isoflurane anesthesia. Muscles were rapidly dissected, blotted dry, weighed, and snap frozen in liquid nitrogen. Muscles for histological analysis were cut transversely at the mid-belly with a scalpel blade, oriented longitudinally on a plastic support, covered with a thin layer of optimal cutting temperature (OCT) compound, and rapidly frozen by plunging into isopentane cooled on liquid nitrogen. Samples were stored at −80°C until further analysis.

### Histological Analysis of Muscle Inflammation and Regeneration

Tissue cross-sections (10 μm) were cut from the muscle mid-belly in a cryostat at −20°C and adhered to SuperFrost Plus slides. Sections were air dried and then stained with hematoxylin and eosin (H & E). Slides for immune cell staining were fixed in acetone for 10 min at −20°C and then air dried. Satellite cell slides were fixed in 4% paraformaldehyde (PFA) for 15 min at room temperature, quenched with hydrogen peroxide, and heat mediated antigen retrieval performed. Unfixed tissue sections were used for muscle fiber type staining as described in detail by Bloemberg et al. 2012 with minor modifications (107). Prepared slides were blocked in 10% normal goat serum (GS, Invitrogen 10000C) or Mouse on Mouse (M.O.M) blocking reagent (Vector Laboratories, MKB-2213) prior to overnight incubation at 4°C with primary antibodies. The following day, slides were incubated with appropriate secondary antibodies and mounted using Fluorescence Mounting Medium (Agilent Dako, S302380). Fluorescent images captured using a Nikon A1 confocal microscope.

### Immunohistochemistry Antibodies

Primary antibodies used on mouse muscle included embryonic myosin heavy chain (DSHB, F1.652s, 1:20), Pax7 (DSHB, Pax7c, 1:100), myosin heavy chain type I (DSHB, BA-D5c, 1:100), myosin heavy chain type IIa (DSHB, SC-71c, 1:100), myosin heavy chain type IIb (DSHB, BF-F3c, 1:100), laminin (Abcam, ab7463, 1:200), rat anti-mouse Ly-6G (Gr-1) (BD Biosciences, BD550291, 1:50), rat anti-mouse CD68 (Bio-Rad, MCA1957, 1:50), and rabbit polyclonal CD163 (Santa Cruz, sc-33560, 1:50). Primary antibodies used on rat muscle included myosin heavy chain type I (DSHB, BA-D5c, 1:100), myosin heavy chain type IIa (DSHB, SC-71c, 1:100), myosin heavy chain type IIb (DSHB, BF-F3c, 1:100), mouse anti-rat granulocytes (HIS-48) (Abcam, Ab33760, 1:20), mouse anti-rat CD68 (Abcam, ab31630, 1:50), and rabbit polyclonal CD163 (Santa Cruz, sc-33560, 1:50). Antibody binding was visualized with standard Alexa Fluor secondary antibodies (Invitrogen, 1:500 in PBS) except for Pax7 which was detected using a Tyramide SuperBoost Kit (Invitrogen, B40913). Fluorescent dyes including 4′,6-diamidino-2-phenylindole (DAPI, Invitrogen, D21490, 2 μg/mL), wheat germ agglutinin (WGA) Alexa Fluor 647 conjugate (Invitrogen, W32466, 5 μg/mL), WGA CF405S conjugate (Biotium, 29027, 100 μg/mL), and phalloidin (Invitrogen, ActinRed 555 ReadyProbes, R37112) were used to counterstain cell nuclei, extracellular matrix, and muscle fibers respectively in particular experiments.

### Image Analysis

Muscle tissue morphology was analyzed on stitched panoramic images of the entire muscle cross-section by high-throughput fully automated image analysis with the MuscleJ plugin for FIJI (108), with few exceptions. At day 5 post-BaCl_2_ injury, regenerating (eMHC^+^) fiber number and size was specifically quantified on the stitched panoramic images of the entire TA muscle cross-section by in-house semi-automated image analysis using ImageJ/FIJI (Supplemental Figure 3). In all cases the cross-sectional area (CSA) measurement of each individual myofiber (technical replicates) were averaged to obtain a single biological replicate for each muscle sample. Immune cells were manually counted throughout the entire mouse TA cross-section and normalized to tissue area as determined by MuscleJ or from five non-overlapping 20 x fields of view captured from within the inflammatory lesion of the rat plantaris (Supplemental Figure 1). In the latter case, the results of the five images were averaged to obtain a single biological replicate for each muscle sample. In all cases, the experimenter was blinded to the experimental group.

### Muscle Force Testing

Mice were anesthetized with 2% isoflurane and placed on a heated platform. The distal half of the TA muscle was isolated by dissecting the overlying skin and fascia. The knee joint was immobilized and a 4–0 silk suture tied around the distal TA tendon which was severed from its boney insertion and tied to the lever arm of a servomotor (6650LR, Cambridge Technology). A saline drip warmed to 37°C was continuously applied to the exposed muscle. The peroneal nerve was stimulated with 0.2 ms pulses using platinum electrodes with the stimulation voltage and muscle length adjusted to obtain optimal muscle length (L_o_) maximum isometric twitch force (P_t_). The TA was then stimulated at increasing frequencies while held at L_o_ until maximum isometric tetanic force (P_o_) was achieved. One-minute rest was allowed between each tetanic contraction. Muscle length then measured with calipers and optimum fiber length (L_f_) determined by multiplying L_o_ by the TA muscle L_f_/L_o_ ratio of 0.6 (109). The cross-sectional area (CSA) of the muscle was calculated by dividing muscle mass by the product of L_f_ and 1.06 mg/mm^3^ (110). Specific P_o_ (sP_o_) was calculated by dividing P_o_ by muscle CSA.

### LC-MS/MS Based Metabolipidomic Profiling of Muscle Tissue

TA muscle samples were mechanically homogenized in 1 mL of 50 mM phosphate, pH 7.4 with 0.9% saline (PBS) using a bead mill with reinforced tubes and zirconium beads (Precellys). The tissue homogenates were centrifuged at 3,000 x g for 5 min and the supernatant was collected. Supernatants (0.85 ml) were spiked with 5 ng each of 15(S)-HETE-d8, 14(15)-EpETrE-d8, Resolvin D2-d5, Leukotriene B4-d4, and Prostaglandin E1-d4 as internal standards (in 150 μl methanol) for recovery and quantitation and mixed thoroughly. The samples were then extracted for polyunsaturated fatty acid metabolites using C18 extraction columns as described earlier (39, 41, 111–113). Briefly, the internal standard spiked samples were applied to conditioned C18 cartridges, washed with 15% methanol in water followed by hexane and then dried under vacuum. The cartridges were eluted with 2 x 0.5 ml methanol with 0.1% formic acid. The eluate was dried under a gentle stream of nitrogen. The residue was re-dissolved in 50 μl methanol-25 mM aqueous ammonium acetate (1:1) and subjected to LC-MS analysis.

HPLC was performed on a Prominence XR system (Shimadzu) using Luna C18 (3μ, 2.1×150 mm) column. The mobile phase consisted of a gradient between A: methanol-water-acetonitrile (10:85:5 v/v) and B: methanol-water-acetonitrile (90:5:5 v/v), both containing 0.1% ammonium acetate. The gradient program with respect to the composition of B was as follows: 0-1 min, 50%; 1-8 min, 50-80%; 8-15 min, 80-95%; and 15-17 min, 95%. The flow rate was 0.2 ml/min. The HPLC eluate was directly introduced to ESI source of QTRAP5500 mass analyzer (ABSCIEX) in the negative ion mode with following conditions: Curtain gas: 35 psi, GS1: 35 psi, GS2: 65 psi, Temperature: 600 °C, Ion Spray Voltage: −1500 V, Collision gas: low, Declustering Potential: −60 V, and Entrance Potential: −7 V. The eluate was monitored by Multiple Reaction Monitoring (MRM) method to detect unique molecular ion – daughter ion combinations for each of the lipid mediators using a scheduled MRM around the expected retention time for each compound. Optimized Collisional Energies (18 – 35 eV) and Collision Cell Exit Potentials (7 – 10 V) were used for each MRM transition. Spectra of each peak detected in the scheduled MRM were recorded using Enhanced Product Ion scan to confirm the structural identity. The data were collected using Analyst 1.7 software and the MRM transition chromatograms were quantitated by MultiQuant software (both from ABSCIEX). The internal standard signals in each chromatogram were used for normalization, recovery, as well as relative quantitation of each analyte.

LC-MS data was analyzed using MetaboAnalyst 4.0 (114). Analytes with >50% missing values were removed from the data set and remaining missing values were replaced with half of the minimum positive value in the original data set. Heat maps were generated using the Pearson distance measure and the Ward clustering algorithm following autoscaling of features (analytes) without data transformation. Targeted statistical analysis was performed on a predetermined subset of metabolites of interest.

### Whole Tissue RNA Extraction and RT-PCR

Muscle was homogenized in TRIzol reagent using a bead mill. RNA was isolated by Phenol/Chloroform extraction and RNA yield determined using a NanoDrop Spectrophotometer (Nanodrop 2000c). Genomic DNA was removed by incubation with DNase I (Ambion, AM2222) followed by its heat inactivation. Total RNA (1 μg) was reversed transcribed to cDNA using SuperScript™ VILO™ Master Mix (Invitrogen, 11-755-050) and RT-PCR performed on a CFX96 Real-Time PCR Detection System (Bio-Rad, 1855195) in duplicate 20 μL reactions of iTaq™ Universal SYBR^®^ Green Supermix (Bio-Rad, #1725124) with 1 μM forward and reverse primers (Table 1). Duplicate technical replicates were averaged to obtain single Ct value from individual muscle sample as a biological replicate. Relative mRNA expression was determined using the 2^−ΔΔCT^ method with *Actb* and *Gapdh* serving as endogenous controls from mouse (BaCl_2_ induced injury) and rat (synergist ablation) experiments respectively. Primer sequences used are listed in Table 1.

**Table 1:**
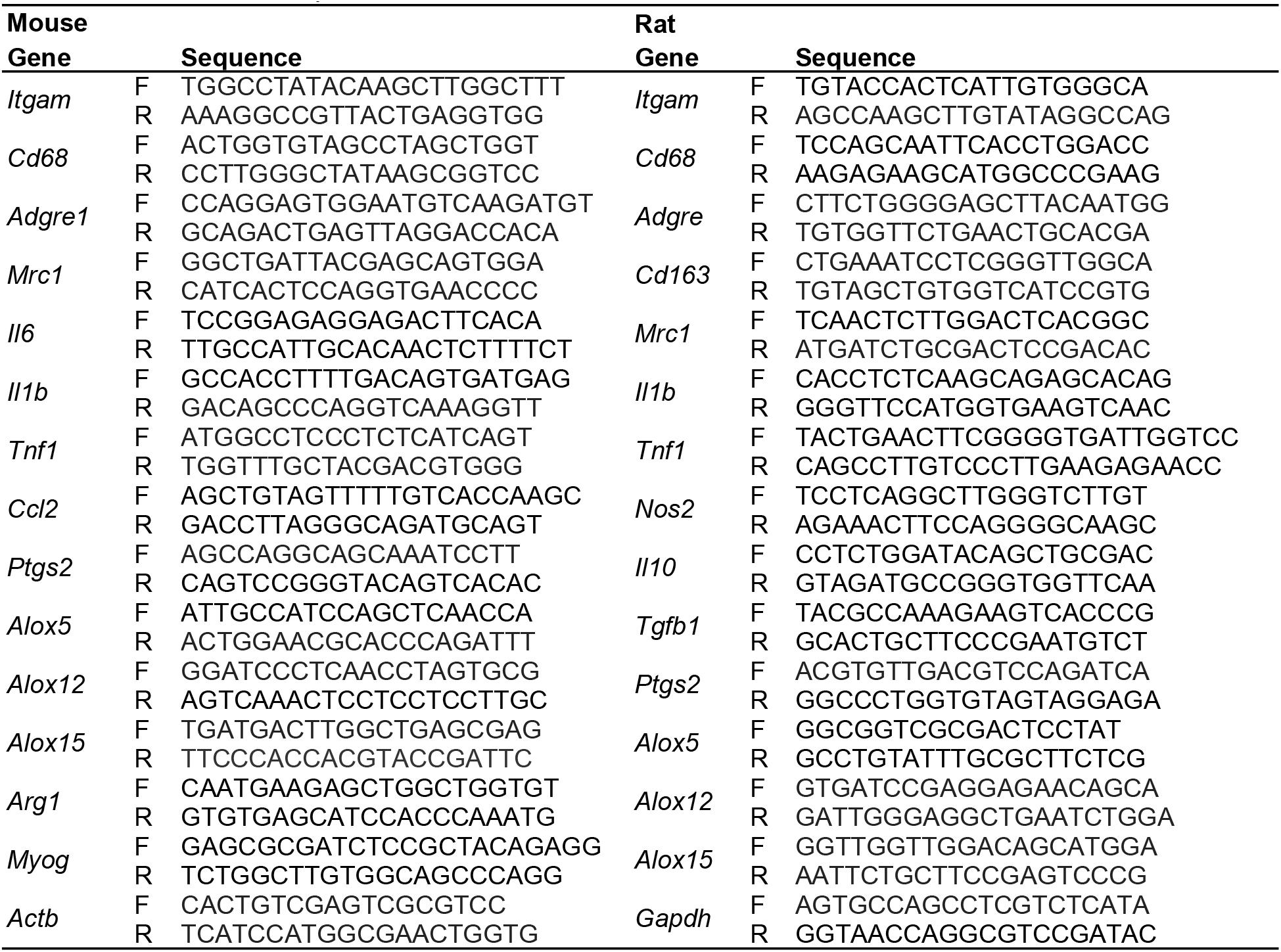
Real-time PCR primers

### Flow Cytometry Analysis of Muscle Inflammation

Both TA muscles from one mouse were pooled, finely minced, and digested for 60 min at 37°C for in Hank’s balanced salt solution lacking calcium and magnesium (HBSS−/−, Fisher Scientific), supplemented with 250 U/mL collagenase II (ThermoFisher), 4.0 U/mL dispase (Sigma-Aldrich, St. Louis, MO), and 2.5 mmol/L CaCl_2_ (Sigma-Aldrich, St. Louis, MO). The resulting digest solution was filtered through a 40-μm cell strainer and centrifuged at 350 x g for 5 min. Cells were Fc blocked with a CD16/CD32 antibody (ThermoFisher, 14-0161-82) for 10 minutes at 4°C and then incubated for 30 min on ice with primary antibodies including CD45-PE, Ly-6G-FITC (1A8) (BD Pharmingen, Franklin Lakes, NJ), CD64-APC (X54-5/7.1), CD11c-APCe780, (Biolegend, San Diego, CA), and Live/Dead fixable violet (ThermoFisher). Flow cytometry was performed using a LSRFortessa cell analyzer (BD Biosciences, San Jose, CA) and data analyzed with FlowJo software.

### FACS Enrichment of Muscle Satellite Cells for RNA-Seq

Both TA muscles from one mouse were pooled, finely minced, and digested for 60 min at 37°C in Dulbecco’s Modified Eagle’s Medium (DMEM) with 0.2% [~5,500 U/mL] collagenase Type II and 2.5 U/mL dispase II. The resulting digest solution was filtered through a 70 μm cell strainer and then centrifuged at 350 x g for 5 min. Cells were resupsended and incubated for 30 min on ice with primary antibodies including Sca-1-APC (Biolegend [Clone D7; 108112]), CD45-AF488 (Biolegend [Clone 30-F11; 103121]), CD11b-APC (Biolegend [Clone M1/70; 101212]), Ter119-APC (Biolegend [Clone TER-119; 116212]), CD29/β1-integrin-PE (Biolegend [Clone HMb-1; 102208]), and CD184/CXCR-4-BIOTIN (BD Biosciences [Lot 6336587; 551968]). Cells were then washed, centrifuged, and re-suspended in PECy7-STREPTAVIDIN secondary antibody (eBioscience [Lot 4290713; 25-4317-82]). The stained cells were filtered through a 35 μm cell strainer, propidium iodide (PI) viability dye added (1 μg) (Thermo Fisher, Waltham, MA), and fluorescence activated cell sorting (FACS) performed using a BD FACSAria III Cell Sorter (BD Biosciences, San Jose, CA). Muscle stem cells (satellite cells, MuSCs), classified here as APC/FITC (Sca-1, CD11b, Ter119, CD45) double-negative and PE/PECy7 (β1-integrin/CXCR-4) double-positive cells, were sorted into Trizol reagent and stored at −80°C.

### RNA Sequencing

RNA was extracted from FACS sorted MuSCs using the miRNeasy Micro Kit (Qiagen, 217084). RNA concentration and integrity were measured with a Nanodrop spectrophotometer (Nanodrop 2000c) and Bioanalyzer High Sensitivity RNA Chip (Agilent 2100). High-quality RNA (10 ng, RIN>9) was used to produce cDNA libraries using SmartSeq v4 (Clontech, 634888). cDNA was prepared into sequencing libraries using 150 pg of full-length cDNA amplicons with the Illumina® Nextera XT DNA Library Preparation Kit with dual index barcodes. Barcoded libraries were pooled and sequenced on an Illumina NextSeq 550 using 76-bp single-ended reads (GEO accession no. https://www.ncbi.nlm.nih.gov/geo/query/acc.cgi?acc=).

### RNA-Seq Data Processing and Analysis

Single-stranded RNAseq data was aligned to the mm10 reference genome with the STAR algorithm (STAR_2.5.0a) and RSEM quantification applied to the aligned reads. Differentially expressed genes were identified in R using Limma-Voom analysis. Expected counts were first Voom transformed to counts per million (CPM) and then surrogate variable analysis was performed with the SVA package. Surrogate variables were quantified and removed from the data matrix. All pairwise contrasts were examined to identify differentially expressed genes between vehicle and RvD1 treated mice. Thresholds of p-adjusted < 0.05 and log_2_-fold-change >=1 were used to call differential genes. Enrichment analysis of gene lists (>2-fold, irrespective of p-value) was assessed using NetworkAnalyst 3.0 (115)

### Muscle Cell Culture

C2C12 myoblasts (ATCC, CRL-1772) were grown at 37°C, 5% CO_2_ in DMEM (Gibco,11995-073), supplemented with 10% fetal bovine serum (FBS, Corning, MT35015CV) and antibiotics (penicillin 100 U/mL, streptomycin 100 μg/mL) (Gibco, 15140122). Myoblasts were plated at a cell density of 2.5 × 10^4^ /cm^2^ in 12-well plates in growth media and allowed to proliferate and crowd for 72 h. Confluent myoblasts were then switched to differentiation media consisting of DMEM (Gibco,11995-073) supplemented with 2% horse serum (HS, Gibco) and antibiotics. Experimental treatments were prepared in differentiation media immediately prior to use and provided to cells either at the onset of myoblast differentiation or to fused myotubes at day 3 post-differentiation. To assess cellular morphology, myotubes were fixed in 4% paraformaldehyde (PFA), permeabilized with 0.2% Triton X-100, and blocked in 1% bovine serum albumin (BSA) prior to overnight incubation at 4°C with a sarcomeric myosin antibody (DSHB, MF-20s, 1:20). The following day, cells were incubated with a Goat Anti-Mouse IgG (H + L) Alexa Fluor 488 conjugated secondary antibody (Invitrogen, A28175) and DAPI to counterstain cell nuclei (Invitrogen, D21490, 2 μg/mL). Stained cells were visualized with a Nikon A1 inverted confocal microscope and fluorescent images captured at 10 x magnification throughout a 5 × 5 field grid encompassing an area of 30 mm^2^ surrounding the center of the culture cell. Six non-consecutive images from the same coordinates within this grid were randomly selected for manually analysis using ImageJ/FIJI. The results of these six images was averaged to obtain a single biological replicate for each culture well.

### Murine Bone Marrow Derived MΦ

Bone marrow was collected from tibias and femurs of 4-6 mo female C57BL/6 mice and cultured for 7 days at 37°C and 5% CO_2_ in DMEM (Gibco,11995-073) supplemented with 10% FBS, antibiotics, and 20 ng/mL recombinant murine GM-CSF (Bio-legend, 576304). Cells were then washed with PBS to remove non-adherent cells and adherent BMMs were detached by incubation in TrypLE Select at 37°C (Gibco, 12563011) followed by gentle cell scraping. BMMs were plated into 96 well plates at 1 × 10^5^ cells/well in growth media lacking GM-CSF and allowed to adhere overnight before use. Primary cells derived from each individual host mouse were considered as biological replicates.

### Human Peripheral Blood Monocyte Derived MΦ

Venous blood was collected from human volunteers by venipuncture into EDTA vacutainers. Peripheral blood mononuclear cells (PBMCs) were isolated by density gradient centrifugation using Lympholyte®-poly Cell Separation Media (Cedarlane, CL5071). Monocytes were then purified from isolated PBMCs by negative selection using a Human Monocyte Isolation Kit (Stem Cell Technologies, 19359) and cultured for 7 days in RPMI 1640 media (Gibco, 11875119) with 10% FBS (Corning, MT35015CV), antibiotics, and human recombinant GM-CSF (20 ng/mL) (R&D Systems, 215-GM-010). Cells then were washed with PBS to remove non-adherent cells and detached by incubation in TrypLE Select at 37°C followed by gentle cell scraping. MΦ were plated in 96 well plates at a density of 5 × 10^4^ cell/well and allowed to adhere overnight. Human MΦ were then polarized to a M1 activation state with 30 ng/mL human recombinant IFN-γ (R&D Systems, 285-IF-100) and LPS (100 ng/mL) for 24 hours prior to use. Primary cells from each human donor were considered to be biological replicates.

### MΦ Phagocytosis

Human or murine MΦ were pre-treated for 15 min with RvD1 (1-100 nM) or vehicle control (0.1% ethanol) in respective growth media which was then replaced with pHrodo Green *E. Coli* Bio Particles (Invitrogen, P35366) prepared in hanks balanced salt solution containing calcium and magnesium (HBSS+/+) and incubated at 37°C for 1 hour in the continued presence of RvD1 treatment. Non-engulfed *E. coli* BioParticles were removed by washing with HBSS+/+ and intracellular fluorescence measured using a plate reader at an excitation/emission of 509/533 nm. Experiments were performed with cells derived from 3-4 donor hosts each in triplicate wells. Triplicates were averaged to obtain a single measurement with donor host considered as biological replicates. Data is presented as the percentage change in fluorescence intensity when compared to vehicle treated MΦ obtained from the same host.

### Statistics

Data is presented as the mean ± SEM with raw data from each individual biological replicate also shown. Statistical analysis was performed in GraphPad Prism 7. Between group differences were tested by two-tailed unpaired students t-tests (2 groups) or by a one-way analysis of variance (ANOVA) followed by pair-wise Holm-Sidak post-hoc tests (≥3 groups). For time-course experiments, multiple comparison testing was made compared to a single baseline control group. p≤0.05 was used to determine statistical significance.

### Study approval

All animal experiments were approved by the University of Michigan Institutional Animal Care and Use committee (IACUC) (PRO00008744 & PRO00006079). Experiments with human participants were approved by the University of Michigan Institutional Review Board (IRBMED) (HUM00158470).

### Data deposition

The RNA-seq data reported in this paper have been deposited in the Gene Expression Omnibus (GEO) database, https://www.ncbi.nlm.nih.gov/geo (accession no. GSE146785 https://www.ncbi.nlm.nih.gov/geo/query/acc.cgi?acc=GSE146785)

## Author contributions

J.F.M and S.V.B conceived the study. S.V.B, K.R.M, P.C.D.M and C.A.A supervised the work. J.F.M, L.A.B, C.A.A, K.B.S, and S.V.B designed the experiments. J.F.M, L.A.B, E.L, C.F, J.L, J.A.C-M, D.C.S, and C.D performed the experiments. J.F.M, L.A.B, J.L, and K.R.M analyzed the data. J.F.M prepared the figures and wrote the manuscript with input from all authors.

## Acknowledgments

This work was supported by the Glenn Foundation for Medical Research Post-Doctoral Fellowship in Aging Research (JFM), the National Institutes of Health (NIH) under the awards R01 (AG050676) (SVB), PO1 (AG051442) (SVB), P30 (AR069620) (CAA, SVB) and S10 (RR027926) (KRM), together with the 3M Foundation (CAA), American Federation for Aging Research (CAA), the University of Michigan Geriatrics Center/National Institute of Aging P30 (AG024824) (CAA, SVB), the University of Michigan Biomedical Engineering Department (CAA), the Department of Defense and Congressionally Directed Medical Research Program W81XWH18SBAA1-1257992 (CAA), and the University of Michigan Department of Orthopedic Surgery. The authors acknowledge expert assistance of Alison C. Ludzki, Suzette Howton and Professor Jeff Horowitz from the Substrate Metabolism Laboratory in obtaining human blood specimens and members of the Benjamin Levi Laboratory for helpful scientific discussions.

## Supplemental Figure Legends

**Supplemental Figure 1.**
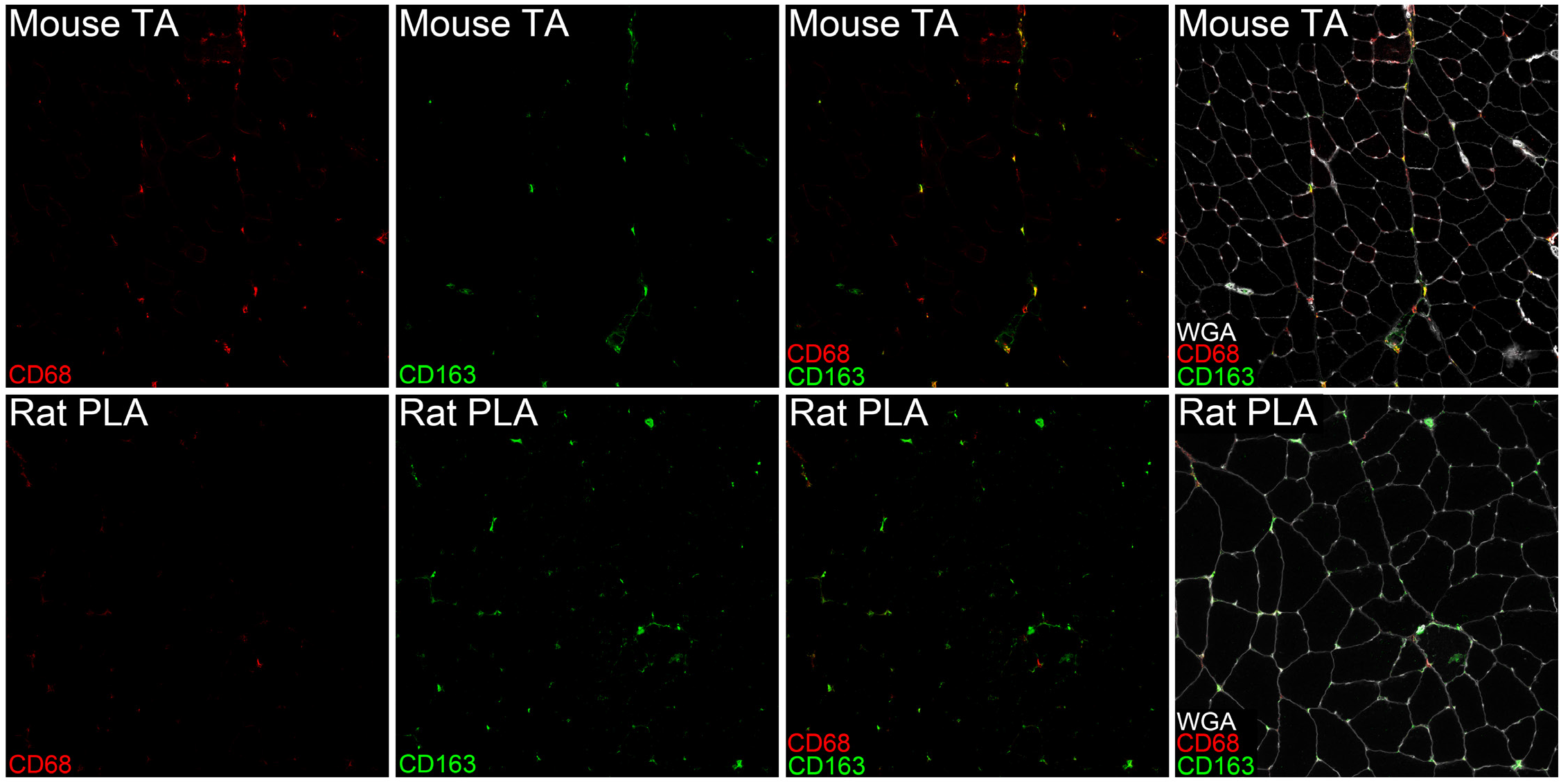
Resident skeletal muscle macrophage population: Cross-sections of uninjured mouse tibialis anterior (TA) and rat plantaris (PLA) muscles were stained with primary antibodies including rat anti-mouse CD68 (Bio-Rad, MCA1957, 1:50) or mouse anti-rat CD68 (ED1) (Abcam, ab31630, 1:50) in combination with rabbit polyclonal CD163 (ED2) (Santa Cruz, sc-33560, 1:50). Uninjured mouse TA muscles contained many resident CD163^+^ cells, the majority of which showed clear co-localization of CD68. In contrast, the many resident CD163^+^ (ED2) cells present in uninjured rat plantaris muscles showed little if any co-expression of the rat analog of CD68 (ED1).

**Supplemental Figure 2.**
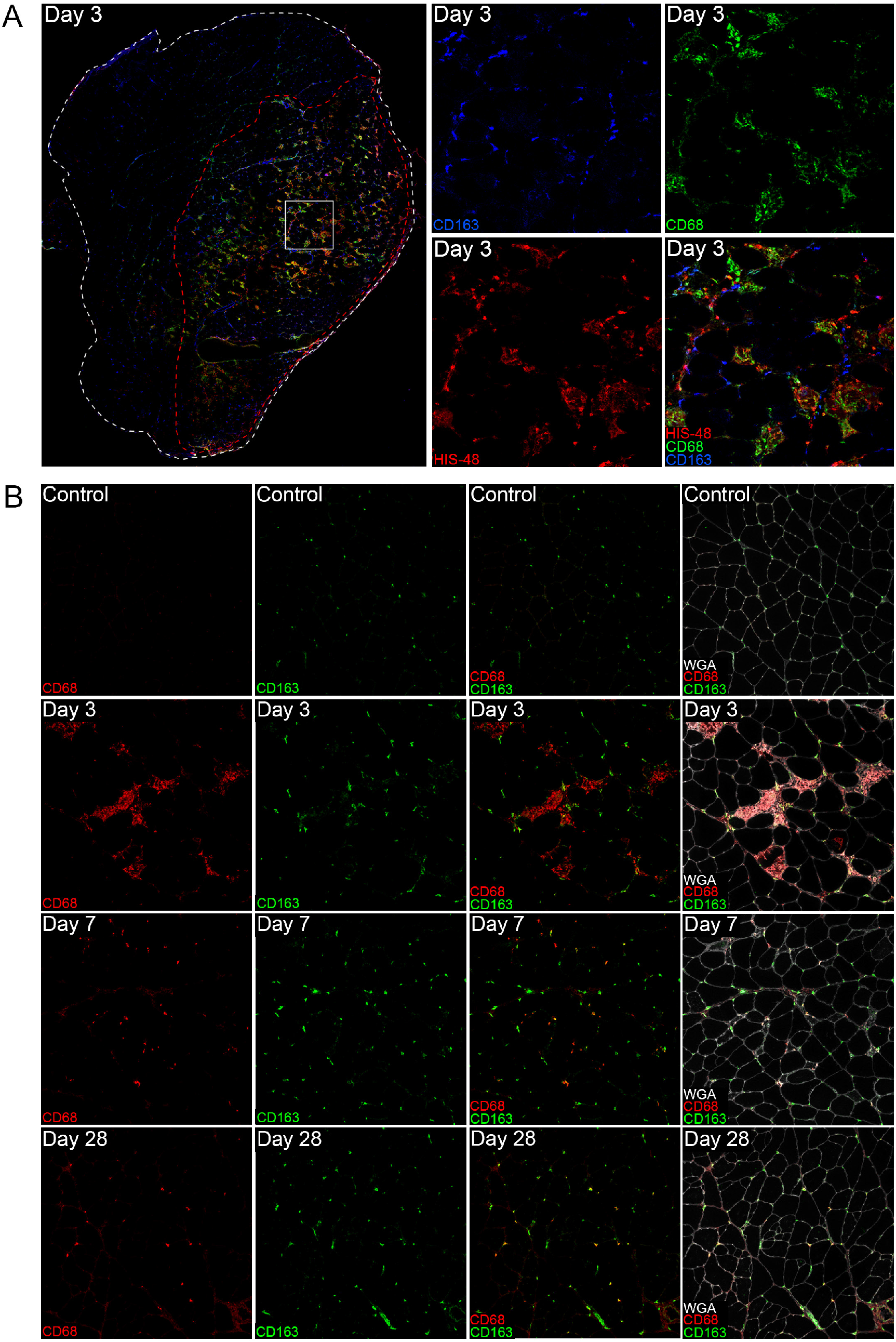
Functional overload induced skeletal muscle inflammation: A: Rat plantaris muscles at day 3 following functional overload induced by synergist ablation surgery were stained with a cocktail of antibodies for polymorphonuclear cells (PMNs, HIS-48), ED1 macrophages (MΦ, CD68), and ED2 MΦ (CD163). At least three distinct innate immune cell types were present within the inflammatory zone (red outline) including PMNs (HIS-48^+^CD68^−^CD163^−^ cells), inflammatory ED1 monocyte/ MΦ (CD68^+^CD163^−^HIS-48^−^cells), and resident-like ED2 MΦ (CD163^+^CD68^−-^HIS-48^−^cells). Many resident CD163^+^ cells, but few CD68^+^ or HIS-48^+^ cells were present outside of the inflammatory zone. B: At day 3 following synergist ablation surgery there was robust infiltration of the overloaded plantaris muscle by many inflammatory CD68^+^CD163^−^ cells (e.g. ED1 MΦ). At day 7 and 28 of synergist ablation smaller numbers of CD68^+^ cells persisted within the overloaded plantaris, many of which showed a clear increase in the co-expression of CD163 (e.g. ED2 MΦ).

**Supplemental Figure 3.**
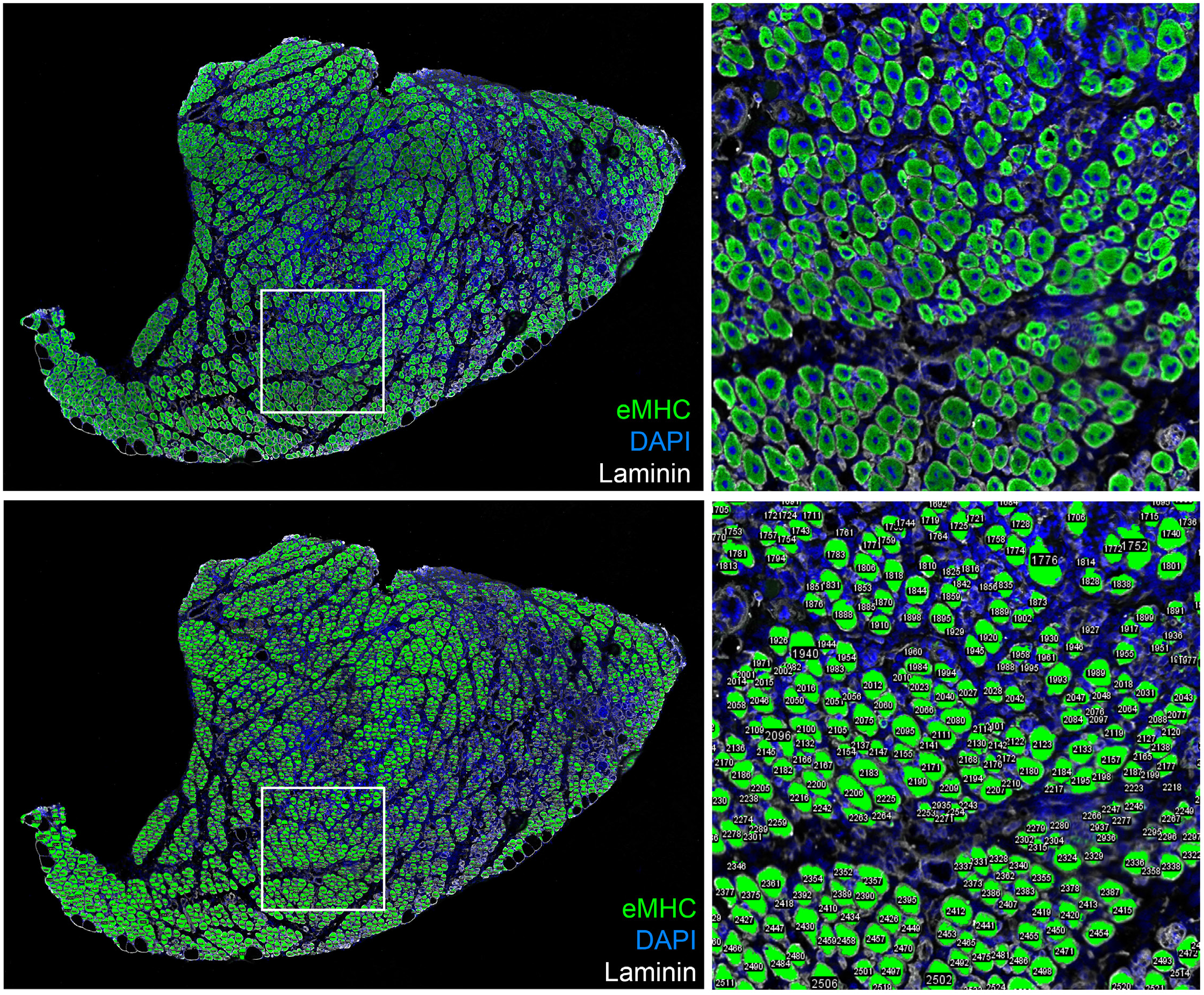
Representative staining and analysis of regenerating myofibers for embryonic myosin heavy chain (eMHC) following skeletal muscle injury: Mouse tibialis anterior (TA) muscles at day 5 following injury induced by intramuscular injection of barium chloride (BaCl_2_) were stained with an antibody against eMHC (DSHB, F1.652s, 1:20). The number and size of each regenerating (eMHC^+^) myofiber were analyzed on stitched panoramic images of the entire TA muscle section via semi-automated analysis using image J/FIJI software. Numeric labels mark those muscle fibers which were counted and measured by the software.

**Supplemental Figure 4.**
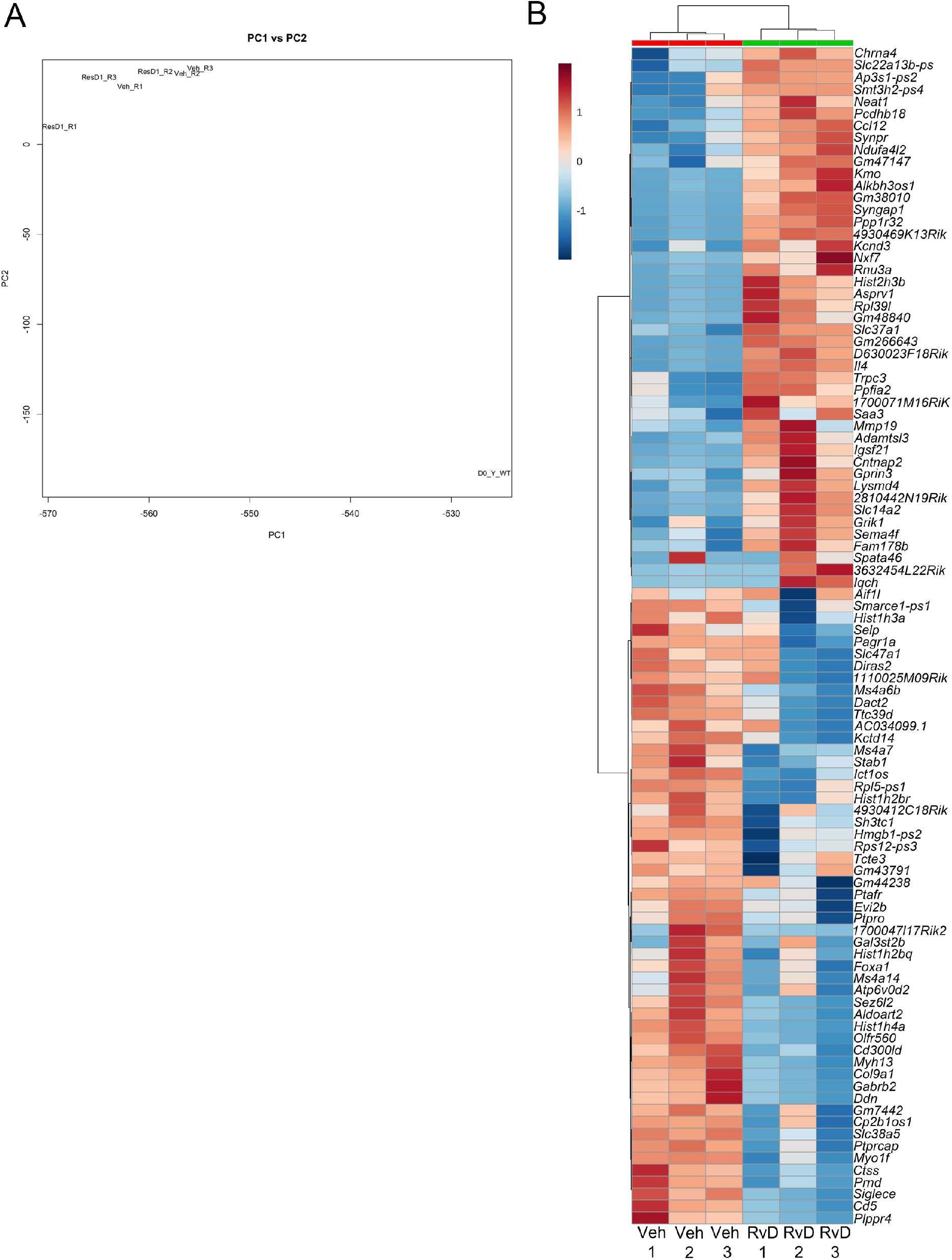
Muscle injury markedly impacts the global muscle satellite cell (MuSC) transcriptome with minimal overall effect of resolvin D1 (RvD1) treatment: A: Multidimensional scaling (MDS) plot showing results of unsupervised principle component analysis (PCA) of RNA sequencing data of the muscle satellite cell (MuSC) transcriptome. MuSCs were isolated from the tibialis anterior (TA) at day 3 following muscle injury induced by intramuscular injection of barium chloride (BaCl_2_) for three biological replicates per group of mice receiving daily intraperitoneal injection with either resolvin D1 (“ResD1_R1-R3”) or vehicle control (“Veh_R1-R3”). The transcriptome of MuSCs isolated from the entire hind-limb musculature of an age and gender matched uninjured control mouse is shown for comparison (D0_Y_WT). B: Top 100 muscle satellite cell (MuSC) genes modulated by resolvin D1 (RvD1) treatment following muscle injury. Heat map displaying the relative gene expression profiles of bulk MuSCs isolated from the tibialis anterior (TA) muscle as determined by RNA-sequencing at day 3 following injury induced by intramuscular injection of 50 μL of 1.2% barium chloride (BaCl_2_) from six mice randomized to receive daily systemic treatment with either RvD1 (100 ng/mouse) or vehicle control (0.1% ethanol).

**Supplemental Figure 5.**
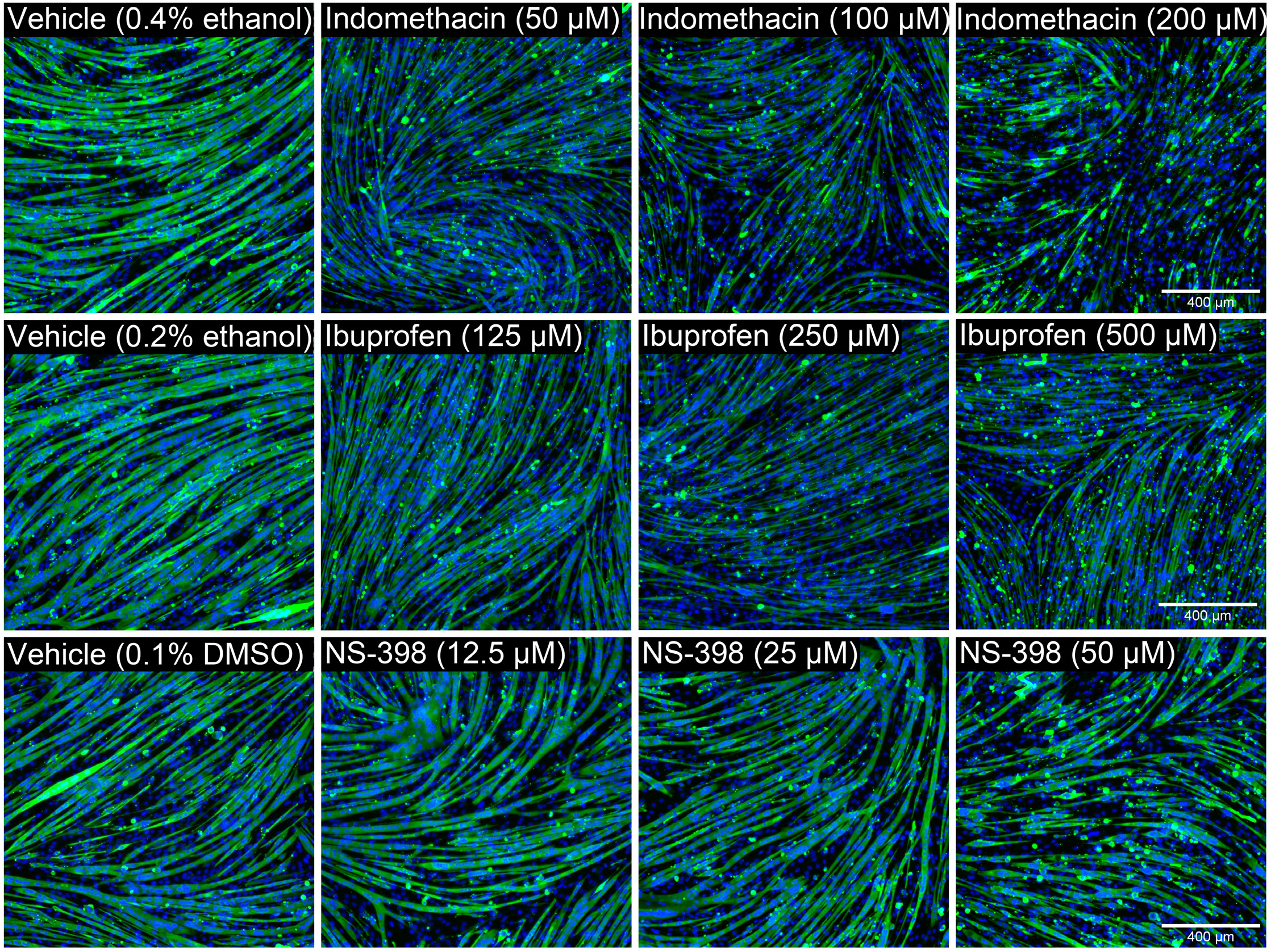
Dose-dependent inhibition of *in-vitro* myogenesis by non-steroidal anti-inflammatory drugs (NSAIDs): Myogenic precursor cells (C2C12 myoblasts) were induced to undergo myogenic differentiation via serum deprivation in the presence of increasing doses of NSAIDs including indomethacin, ibuprofen, and NS-398. At 3-days post-differentiation, the resulting fused myotube cultures were fixed in 4% paraformaldehyde and stained with an antibody against sarcromeric myosin. Cell nuclei were counterstained with DAPI and cells were visualized by fluorescence microscopy.

**Supplemental Tables 1A:** Cyclooxygenase metabolite concentration (pg/mg) in the mouse tibialis anterior (TA) muscle following BaCl_2_ injury.

| Pathway | Substrate | Analyte | Sham |  |  | BaCl<sub>2</sub> |  |  |
| --- | --- | --- | --- | --- | --- | --- | --- | --- |
|  |  |  | 1D | 3D | 5D | 1D | 3D | 5D |
| Cyclooxygenase | 20:3n-6 | PGE1 | ND | ND | ND | ND | ND | 2.49 ± 1.10 |
|  |  | PGF1alpha | ND | ND | ND | ND | ND | ND |
|  |  | 15-keto PGE1 | ND | ND | ND | ND | ND | ND |
|  |  | 13,14dhPGE1 | ND | ND | ND | ND | ND | ND |
|  |  | 13,14dh-15k-PGE1 | ND | ND | ND | ND | ND | ND |
|  |  | D17-PGE1 | ND | ND | ND | ND | ND | ND |
|  |  | 15(R)-PGE1 | ND | ND | ND | ND | ND | ND |
|  |  | Bicyclo PGE1 | ND | ND | ND | ND | ND | ND |
|  |  | 19(R)-hydroxy PGE1 | ND | ND | ND | ND | ND | ND |
|  |  | 2,3-dinor PGE1 | ND | ND | ND | ND | ND | ND |
|  |  | 6-keto PGE1 | ND | ND | ND | ND | ND | ND |
|  | 20:4n-6 | TXB2 | 9.08 ± 1.53 | 8.33 ± 1.03 | 7.09 ± 0.83 | 5.10 ± 0.32 | 27.14 ± 10.53 | 13.60 ± 3.89 |
|  |  | 12-HHTre | 2.95 ± 0.34 | 2.47 ± 0.21 | 3.20 ± 0.30 | 2.15 ± 0.31 | 6.64 ± 2.15 | 7.00 ± 2.13 |
|  |  | 11dh-2,3-dinor TXB2 | 2.52 ± 0.26 | 3.67 ± 0.48 | 5.48 ± 0.97 | 3.07 ± 0.27 | 3.56 ± 0.78 | 0.79 ± 0.48 |
|  |  | PGA2 | 4.66 ± 0.87 | 5.40 ± 1.53 | 4.52 ± 0.62 | 4.10 ± 0.41 | 7.47 ± 2.42 | 7.00 ± 1.23 |
|  |  | PGD2 | 10.61 ± 4.86 | 27.78 ± 15.50 | 5.40 ± 0.66 | 7.46 ± 3.45 | 60.63 ± 32.01 | 30.41 ± 11.60 |
|  |  | PGE2 | 31.91 ± 11.47 | 53.38 ± 17.51 | 27.63 ± 3.62 | 32.56 ± 8.46 | 69.11 ± 25.47 | 56.78 ± 13.64 |
|  |  | PGF2alpha | 3.93 ± 0.32 | 4.73 ± 0.78 | 4.87 ± 0.22 | 3.20 ± 1.07 | 5.72 ± 0.98 | 7.67 ± 1.68 |
|  |  | 6kPGF1alpha | 3.55 ± 0.49 | 4.59 ± 0.84 | 4.51 ± 1.54 | 4.26 ± 0.82 | 10.43 ± 4.93 | 16.45 ± 5.06 |
|  |  | 15-keto PGE2 | 0.68 ± 0.28 | 1.24 ± 0.33 | 1.01 ± 0.34 | 1.29 ± 0.09 | 2.45 ± 0.79 | 2.29 ± 0.88 |
|  |  | 15-keto PGF2alpha | 1.75 ± 0.75 | 3.30 ± 1.04 | 1.72 ± 0.17 | 2.04 ± 0.42 | 3.82 ± 1.29 | 3.64 ± 0.91 |
|  |  | 13,14dh-15k-PGE2 | 2.08 ± 0.96 | 3.50 ± 1.46 | 2.01 ± 0.38 | 2.09 ± 0.64 | 4.74 ± 1.76 | 3.92 ± 1.09 |
|  |  | tetranor PGEM | ND | ND | ND | ND | ND | 0.22 ± 0.12 |
|  |  | PGJ2 | 1.62 ± 0.30 | 2.74 ± 0.64 | 1.63 ± 0.69 | 1.44 ± 0.37 | 4.39 ± 1.75 | 3.56 ± 0.94 |
|  |  | D12-PGJ2 | 0.98 ± 0.52 | 1.58 ± 0.68 | 0.28 ± 0.26 | 0.76 ± 0.46 | 1.78 ± 0.93 | 1.21 ± 0.74 |
|  |  | 13,14dh-15k-PGD2 | 1.75 ± 0.61 | 3.27 ± 0.92 | 1.63 ± 0.63 | 1.56 ± 0.66 | 4.69 ± 2.51 | 4.24 ± 1.64 |
|  |  | 11dh-TXB2 | ND | ND | ND | ND | ND | ND |
|  |  | 2,3-dinor TXB2 | ND | ND | ND | ND | ND | ND |
|  |  | 13,14dh-15k-PGF2alpha | ND | ND | ND | ND | ND | ND |
|  |  | 8-isoPGF2alpha & 11bPGF2alpha | ND | ND | ND | ND | ND | ND |
|  |  | 19(R)-OH PGF2alpha & 20-OH PGF2alpha | ND | ND | ND | ND | ND | ND |
|  |  | Bicyclo PGE2 | ND | ND | ND | ND | ND | ND |
|  |  | 19(R)-OH PGE2 & 20-OH PGE2 | ND | ND | ND | ND | ND | ND |
|  |  | 15d-D12,14-PGJ2 | ND | ND | ND | ND | ND | ND |
|  |  | 6,15-diketo PGFalpha | ND | ND | ND | ND | ND | ND |
|  |  | iPF-VI | ND | ND | ND | ND | ND | ND |
|  | 20:5n-3 | TXB3 | ND | ND | ND | ND | 2.28 ± 0.93 | ND |
|  |  | 11dh TXB3 | ND | ND | ND | ND | ND | ND |
|  |  | PGD3 | ND | ND | ND | ND | ND | ND |
|  |  | PGE3 | ND | ND | ND | ND | ND | ND |
|  |  | PGF3alpha | ND | ND | ND | ND | ND | ND |
|  |  | 15d-D12,14-PGJ3 | ND | ND | ND | ND | ND | ND |

Values are mean ± SEM of 5 mice/group. ND = Below limits of detection of the assay.

**Supplemental Tables 1B:** Lipoxygenase metabolite concentration (pg/mg) in the mouse tibialis anterior (TA) muscle following BaCl_2_ injury.

| Pathway | Substrate | Analyte | Sham |  |  | BaCl<sub>2</sub> |  |  |
| --- | --- | --- | --- | --- | --- | --- | --- | --- |
|  |  |  | 1D | 3D | 5D | 1D | 3D | 5D |
| Lipoxygenase | 18:2n-6 | 9-HODE | 17.05 ± 3.04 | 18.93 ± 2.70 | 23.32 ± 2.05 | 19.79 ± 1.28 | 47.84 ± 12.90 | 47.88 ± 8.53 |
|  |  | 9-OxoODE | 21.28 ± 3.65 | 23.81 ± 4.40 | 27.24 ± 1.40 | 25.28 ± 2.09 | 54.09 ± 14.09 | 45.54 ± 15.21 |
|  |  | 13-HODE | 124.43 ± 19.14 | 129.16 ± 15.00 | 157.85 ± 17.91 | 124.45 ± 6.83 | 381.38 ± 111.39 | 312.06 ± 82.64 |
|  |  | 13-OxoODE | 37.07 ± 5.09 | 22.43 ± 3.71 | 54.17 ± 2.83 | 46.29 ± 6.46 | 98.27 ± 22.54 | 101.00 ± 32.05 |
|  | 18:3n-3 | 9-HOTrE | 1.09 ± 0.10 | 1.12 ± 0.34 | 1.83 ± 0.20 | 1.23 ± 0.18 | 2.76 ± 0.66 | 2.84 ± 0.93 |
|  |  | 9-OxoOTrE | 0.80 ± 0.05 | 0.95 ± 0.17 | 1.35 ± 0.09 | 0.80 ± 0.09 | 2.05 ± 0.60 | 2.28 ± 0.78 |
|  |  | 13-HOTrE | 2.19 ± 0.61 | 2.03 ± 0.12 | 3.15 ± 0.83 | 1.51 ± 0.20 | 8.98 ± 3.09 | 7.94 ± 3.60 |
|  | 18:3n-6 | 13-HOTrE(gamma) | ND | ND | ND | ND | ND | ND |
|  | 20:2n-6 | 11-HEDE | 0.22 ± 0.14 | 0.23 ± 0.15 | 0.17 ± 0.16 | 0.25 ± 0.14 | 0.51 ± 0.39 | 0.13 ± 0.12 |
|  |  | 15-HEDE | 0.15 ± 0.09 | 0.18 ± 0.07 | 0.17 ± 0.09 | 0.20 ± 0.08 | 1.14 ± 0.45 | 0.68 ± 0.33 |
|  |  | 15-OxoEDE | ND | ND | ND | ND | ND | ND |
|  | 20:3n-9 | 5-HETrE | 0.10 ± 0.03 | 0.17 ± 0.02 | 0.27 ± 0.11 | 0.12 ± 0.01 | 0.30 ± 0.05 | 0.44 ± 0.12 |
|  |  | 8-HETrE | 0.86 ± 0.20 | 0.95 ± 0.14 | 0.84 ± 0.20 | 0.88 ± 0.07 | 2.01 ± 0.55 | 1.93 ± 0.42 |
|  | 20:4n-6 | 5-HETE | 5.70 ± 1.45 | 5.73 ± 0.89 | 7.91 ± 2.22 | 4.05 ± 0.13 | 13.23 ± 3.74 | 25.38 ± 7.64 |
|  |  | 5-oxoETE | 1.14 ± 0.23 | 1.64 ± 0.10 | 1.43 ± 0.09 | 1.95 ± 0.17 | 3.39 ± 0.73 | 4.15 ± 1.26 |
|  |  | 8-HETE | 3.01 ± 0.78 | 2.31 ± 0.32 | 2.52 ± 0.43 | 2.49 ± 0.29 | 6.49 ± 1.72 | 7.55 ± 1.64 |
|  |  | 9-HETE | ND | ND | ND | ND | ND | ND |
|  |  | 11-HETE | 12.37 ± 1.67 | 14.45 ± 2.10 | 10.88 ± 0.53 | 12.99 ± 1.39 | 39.39 ± 12.54 | 41.83 ± 9.49 |
|  |  | 12-HETE | 338.65 ± 89.86 | 230.63 ± 55.65 | 124.93 ± 4.48 | 177.23 ± 30.88 | 930.19 ± 324.22 | 645.96 ± 278.41 |
|  |  | 12-OxoETE | 0.98 ± 0.46 | 0.38 ± 0.17 | 0.58 ± 0.22 | 0.78 ± 0.27 | 2.91 ± 1.15 | 1.55 ± 0.97 |
|  |  | tetranor 12-HETE | 0.24 ± 0.11 | 0.44 ± 0.16 | 0.34 ± 0.17 | 0.05 ± 0.04 | 0.40 ± 0.16 | 0.45 ± 0.08 |
|  |  | 15-HETE | 7.71 ± 2.13 | 7.35 ± 0.67 | 7.54 ± 0.54 | 7.21 ± 0.87 | 24.77 ± 7.71 | 22.17 ± 5.93 |
|  |  | 15-OxoETE | ND | ND | ND | ND | ND | ND |
|  |  | 20-HETE | 4.03 ± 0.65 | 4.11 ± 0.81 | 3.11 ± 0.35 | 3.50 ± 1.00 | 8.86 ± 1.87 | 7.13 ± 1.61 |
|  |  | 5(S),12(S)-DiHETE | 11.86 ± 4.64 | 4.17 ± 0.71 | 3.22 ± 0.26 | 3.66 ± 1.08 | 25.92 ± 11.56 | 24.75 ± 13.75 |
|  |  | 5(S),15(S)-DiHETE | ND | ND | ND | ND | ND | 0.43 ± 0.31 |
|  |  | 8(S),15(S)-DiHETE | ND | ND | ND | ND | ND | ND |
|  |  | LTB4 | ND | ND | ND | ND | ND | ND |
|  |  | 5(S),6(S)-DiHETE | ND | ND | ND | ND | ND | ND |
|  |  | 5,6-DiHETE(n-3)) | ND | ND | ND | ND | ND | ND |
|  |  | 12-OxoLTB4 | ND | ND | ND | ND | ND | ND |
|  |  | 20-hydroxy LTB4 | ND | ND | ND | ND | ND | ND |
|  |  | 20-COOH LTB4 | ND | ND | ND | ND | ND | ND |
|  |  | 18-carboxy dinor LTB4 | ND | ND | ND | ND | ND | ND |
|  | 20:5n-3 | 5-HEPE | 1.07 ± 0.23 | 1.24 ± 0.23 | 1.32 ± 0.15 | 0.83 ± 0.15 | 2.46 ± 0.78 | 3.16 ± 0.84 |
|  |  | 8-HEPE | ND ± ND | ND ± ND | ND ± ND | ND ± ND | 0.64 ± 0.39 | 0.83 ± 0.51 |
|  |  | 9-HEPE | 1.85 ± 0.92 | 0.78 ± 0.48 | 0.35 ± 0.33 | 0.92 ± 0.42 | 7.93 ± 3.14 | 5.30 ± 2.84 |
|  |  | 11-HEPE | 0.60 ± 0.30 | 0.55 ± 0.28 | 0.21 ± 0.19 | 0.56 ± 0.24 | 2.18 ± 0.70 | 2.59 ± 0.52 |
|  |  | 12-HEPE | 90.52 ± 17.66 | 59.47 ± 8.47 | 38.34 ± 3.72 | 36.66 ± 6.68 | 241.63 ± 80.93 | 165.77 ± 75.28 |
|  |  | 15-HEPE | 2.93 ± 1.49 | 2.50 ± 0.39 | 2.86 ± 1.24 | 2.03 ± 0.55 | 12.23 ± 5.19 | 7.73 ± 4.29 |
|  |  | 18-HEPE | 0.68 ± 0.32 | 0.80 ± 0.24 | 1.31 ± 0.12 | 0.95 ± 0.08 | 1.78 ± 0.37 | 2.33 ± 0.39 |
|  |  | LTB5 | ND | ND | ND | ND | 0.47 ± 0.28 | ND |
|  | 22:6n-3 | 4-HDoHE | 4.52 ± 0.64 | 4.82 ± 0.65 | 4.32 ± 0.30 | 4.82 ± 0.58 | 8.63 ± 1.86 | 9.18 ± 1.79 |
|  |  | 7-HDoHE | 1.15 ± 0.11 | 1.22 ± 0.11 | 1.08 ± 0.09 | 1.04 ± 0.32 | 2.37 ± 0.59 | 2.90 ± 0.29 |
|  |  | 8-HDoHE | 2.59 ± 0.29 | 2.34 ± 0.30 | 2.14 ± 0.20 | 2.50 ± 0.44 | 5.51 ± 1.46 | 4.70 ± 0.64 |
|  |  | 10-HDoHE | 5.21 ± 0.99 | 4.41 ± 0.22 | 3.55 ± 0.41 | 5.47 ± 0.69 | 12.98 ± 4.26 | 9.72 ± 2.45 |
|  |  | 11-HDoHE | 3.14 ± 0.99 | 2.71 ± 0.65 | 2.20 ± 0.25 | 2.65 ± 0.39 | 2.04 ± 1.06 | 3.89 ± 1.24 |
|  |  | 13-HDoHE | 3.76 ± 0.47 | 4.10 ± 0.41 | 3.59 ± 0.26 | 5.18 ± 0.85 | 11.39 ± 3.71 | 9.47 ± 1.55 |
|  |  | 14-HDoHE | 48.10 ± 10.93 | 31.80 ± 7.08 | 19.39 ± 3.50 | 36.35 ± 6.49 | 157.17 ± 65.68 | 76.54 ± 37.15 |
|  |  | 16-HDoHE | 4.19 ± 0.48 | 4.12 ± 0.61 | 4.40 ± 0.58 | 4.42 ± 0.46 | 8.82 ± 1.99 | 9.47 ± 1.10 |
|  |  | 17-HDoHE | 3.85 ± 1.66 | 3.35 ± 0.36 | 2.44 ± 0.86 | 3.79 ± 1.13 | 16.25 ± 7.07 | 8.61 ± 4.10 |
|  |  | 20-HDoHE | 2.77 ± 0.74 | 1.81 ± 1.10 | 3.88 ± 0.31 | 4.45 ± 0.61 | 6.81 ± 1.70 | 7.19 ± 1.19 |

Values are mean ± SEM of 5 mice/group. ND = Below limits of detection of the assay.

**Supplemental Tables 1C:** Epoxygenase metabolite concentration (pg/mg) in the mouse tibialis anterior (TA) muscle following BaCl_2_ injury.

| Pathway | Substrate | Analyte | Sham |  |  | BaCl<sub>2</sub> |  |  |
| --- | --- | --- | --- | --- | --- | --- | --- | --- |
|  |  |  | 1D | 3D | 5D | 1D | 3D | 5D |
| Epoxygenase | 18:2n-6 | 9(10)-EpOME | 311.33 ± 22.27 | 364.51 ± 65.00 | 409.82 ± 54.21 | 274.69 ± 64.10 | 675.21 ± 193.81 | 454.83 ± 87.73 |
|  |  | 12(13)-EpOME | 225.24 ± 20.36 | 253.95 ± 32.03 | 312.30 ± 37.50 | 227.99 ± 51.89 | 411.64 ± 102.42 | 275.60 ± 45.32 |
|  |  | 9,10-DiHOME | 10.47 ± 1.00 | 10.58 ± 0.87 | 15.98 ± 2.27 | 12.70 ± 1.49 | 16.29 ± 2.32 | 17.55 ± 2.65 |
|  |  | 12,13-DiHOME | 12.16 ± 1.35 | 11.08 ± 0.97 | 17.86 ± 2.50 | 15.49 ± 2.05 | 19.33 ± 2.90 | 19.14 ± 2.32 |
|  | 20:4n-6 | 5(6)-EpETrE | 1.53 ± 0.29 | 1.50 ± 0.18 | 1.21 ± 0.23 | 1.33 ± 0.15 | 2.63 ± 0.42 | 3.06 ± 0.81 |
|  |  | 8(9)-EpETrE | ND | ND | ND | ND | 4.66 ± 1.14 | 2.75 ± 2.12 |
|  |  | 11(12)-EpETrE | 17.34 ± 1.38 | 18.92 ± 2.64 | 16.60 ± 1.80 | 15.45 ± 2.06 | 36.47 ± 6.19 | 33.05 ± 7.36 |
|  |  | 14(15)-EpETrE | 8.91 ± 0.84 | 9.96 ± 1.34 | 9.52 ± 1.04 | 7.88 ± 1.08 | 19.59 ± 3.12 | 16.32 ± 3.18 |
|  |  | 5,6-DiHETrE | ND | ND | ND | ND | ND | ND |
|  |  | 8,9-DiHETrE | ND | ND | ND | ND | ND | ND |
|  |  | 11,12-DiHETrE | 0.71 ± 0.31 | 1.15 ± 0.17 | 1.56 ± 0.17 | 1.38 ± 0.12 | 2.28 ± 0.44 | 1.74 ± 0.48 |
|  |  | 14,15-DiHETrE | 1.69 ± 0.10 | 1.84 ± 0.19 | 2.73 ± 0.41 | 1.99 ± 0.19 | 3.55 ± 0.47 | 2.89 ± 0.19 |
|  | 20:5n-3 | 8(9)-EpETE | ND | ND | ND | ND | ND | ND |
|  |  | 11(12)-EpETE | ND | ND | ND | ND | ND | ND |
|  |  | 14(15)-EpETE | ND | ND | ND | ND | ND | 1.17 ± 0.74 |
|  |  | 17(18)-EpETE | 1.95 ± 0.21 | 1.58 ± 0.45 | 2.01 ± 0.80 | 1.16 ± 0.29 | 3.32 ± 1.03 | 3.17 ± 0.92 |
|  | 22:6n-3 | 7(8)-EpDPE | 3.75 ± 0.21 | 4.68 ± 1.01 | 2.95 ± 0.31 | 3.08 ± 0.44 | 7.27 ± 1.08 | 5.97 ± 1.11 |
|  |  | 10(11)-EpDPE | 23.55 ± 1.68 | 21.63 ± 3.28 | 20.83 ± 1.95 | 18.67 ± 3.50 | 41.73 ± 4.44 | 29.16 ± 4.90 |
|  |  | 13(14)-EpDPE | 12.58 ± 0.65 | 12.99 ± 1.96 | 11.53 ± 1.14 | 11.29 ± 1.68 | 22.63 ± 3.21 | 16.47 ± 2.26 |
|  |  | 16(17)-EpDPE | 9.18 ± 0.39 | 9.51 ± 1.51 | 8.13 ± 0.80 | 7.83 ± 1.40 | 15.11 ± 1.66 | 10.46 ± 1.77 |
|  |  | 19(20)-EpDPE | 3.95 ± 2.44 | 3.27 ± 3.25 | 6.04 ± 3.79 | 10.77 ± 3.56 | 24.31 ± 4.34 | 15.72 ± 3.18 |
|  |  | 19,20-DiHDoPE | 3.07 ± 0.91 | 2.46 ± 0.22 | 5.13 ± 2.17 | 11.79 ± 5.02 | 14.85 ± 5.69 | 6.42 ± 1.39 |

Values are mean ± SEM of 5 mice/group. ND = Below limits of detection of the assay.

**Supplemental Tables 1D:**
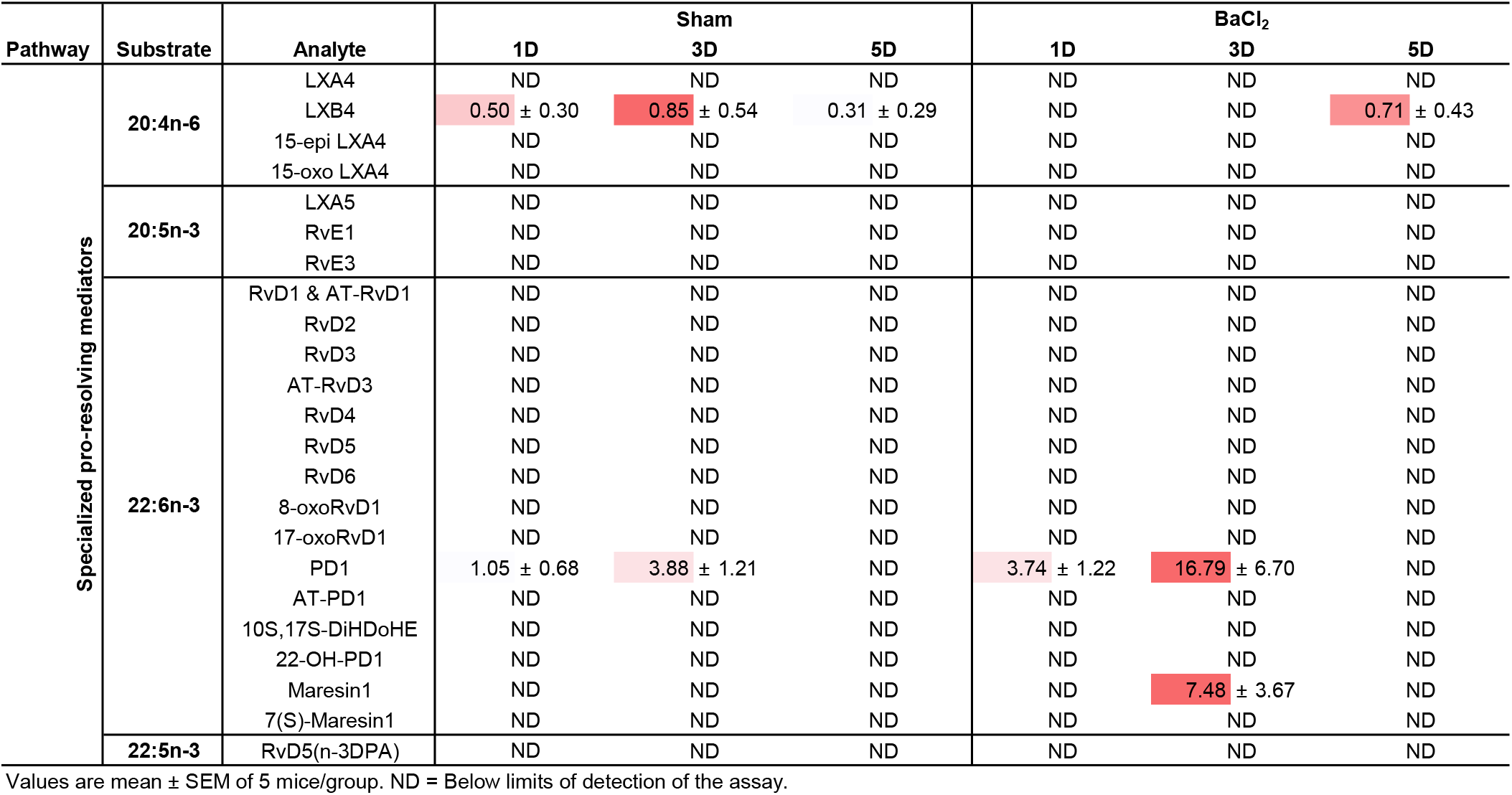
Specialized pro-resolving mediators (pg/mg) in the mouse tibialis anterior (TA) muscle following BaCl_2_ injury.

| Pathway | Substrate | Analyte | Sham |  |  | BaCl<sub>2</sub> |  |  |
| --- | --- | --- | --- | --- | --- | --- | --- | --- |
|  |  |  | 1D | 3D | 5D | 1D | 3D | 5D |
| Specialized pro-resolving mediators | 20:4n-6 | LXA4 | ND | ND | ND | ND | ND | ND |
|  |  | LXB4 | 0.50 ± 0.30 | 0.85 ± 0.54 | 0.31 ± 0.29 | ND | ND | 0.71 ± 0.43 |
|  |  | 15-epi LXA4 | ND | ND | ND | ND | ND | ND |
|  |  | 15-oxo LXA4 | ND | ND | ND | ND | ND | ND |
|  | 20:5n-3 | LXA5 | ND | ND | ND | ND | ND | ND |
|  |  | RvE1 | ND | ND | ND | ND | ND | ND |
|  |  | RvE3 | ND | ND | ND | ND | ND | ND |
|  | 22:6n-3 | RvD1 & AT-RvD1 | ND | ND | ND | ND | ND | ND |
|  |  | RvD2 | ND | ND | ND | ND | ND | ND |
|  |  | RvD3 | ND | ND | ND | ND | ND | ND |
|  |  | AT-RvD3 | ND | ND | ND | ND | ND | ND |
|  |  | RvD4 | ND | ND | ND | ND | ND | ND |
|  |  | RvD5 | ND | ND | ND | ND | ND | ND |
|  |  | RvD6 | ND | ND | ND | ND | ND | ND |
|  |  | 8-oxoRvD1 | ND | ND | ND | ND | ND | ND |
|  |  | 17-oxoRvD1 | ND | ND | ND | ND | ND | ND |
|  |  | PD1 | 1.05 ± 0.68 | 3.88 ± 1.21 | ND | 3.74 ± 1.22 | 16.79 ± 6.70 | ND |
|  |  | AT-PD1 | ND | ND | ND | ND | ND | ND |
|  |  | 10S,17S-DiHDoHE | ND | ND | ND | ND | ND | ND |
|  |  | 22-OH-PD1 | ND | ND | ND | ND | ND | ND |
|  |  | Maresin1 | ND | ND | ND | ND | 7.48 ± 3.67 | ND |
|  |  | 7(S)-Maresin1 | ND | ND | ND | ND | ND | ND |
|  | 22:5n-3 | RvD5(n-3DPA) | ND | ND | ND | ND | ND | ND |

Values are mean ± SEM of 5 mice/group. ND = Below limits of detection of the assay.

**Supplemental Tables 1E:** Cyclooxygenase metabolite concentration (pg/mg) in the rat plantaris muscle in response to functional overload.

| Pathway | Substrate | Analyte | Synergist ablation time-point |  |  |  |
| --- | --- | --- | --- | --- | --- | --- |
|  |  |  | Control | 3D | 7D | 28D |
| Cyclooxygenase | 20:3n-6 | PGE1 | ND | ND | ND | ND |
|  |  | PGF1alpha | ND | ND | ND | ND |
|  |  | 15-keto PGE1 | ND | ND | ND | ND |
|  |  | 13,14dhPGE1 | ND | ND | ND | ND |
|  |  | 13,14dh-15k-PGE1 | ND | ND | ND | ND |
|  |  | D17-PGE1 | ND | ND | ND | ND |
|  |  | 15(R)-PGE1 | ND | ND | ND | ND |
|  |  | Bicyclo PGE1 | ND | ND | ND | ND |
|  |  | 19(R)-hydroxy PGE1 | ND | ND | ND | ND |
|  |  | 2,3-dinor PGE1 | 0.24 ± 0.12 | 0.08 ± 0.04 | ND | ND |
|  |  | 6-keto PGE1 | ND | ND | ND | ND |
|  | 20:4n-6 | TXB2 | 5.54 ± 1.47 | 17.83 ± 5.76 | 5.21 ± 0.92 | 7.83 ± 2.67 |
|  |  | 12-HHTrE | 0.38 ± 0.11 | 1.45 ± 0.50 | 0.27 ± 0.10 | 0.69 ± 0.14 |
|  |  | 11dh-2,3-dinor TXB2 | 1.83 ± 0.19 | 1.38 ± 0.13 | 1.75 ± 0.09 | 1.64 ± 0.17 |
|  |  | PGA2 | 1.52 ± 0.48 | 0.87 ± 0.33 | 0.86 ± 0.60 | 1.03 ± 0.42 |
|  |  | PGD2 | 1.79 ± 0.53 | 3.99 ± 1.89 | 3.04 ± 0.97 | 2.95 ± 0.85 |
|  |  | PGE2 | 2.66 ± 0.52 | 6.52 ± 1.23 | 4.57 ± 0.58 | 4.71 ± 0.93 |
|  |  | PGF2alpha | ND | 0.46 ± 0.23 | 0.24 ± 0.16 | 0.22 ± 0.11 |
|  |  | 6kPGF1alpha | ND | 1.64 ± 0.18 | 0.69 ± 0.20 | 0.36 ± 0.18 |
|  |  | 15-keto PGE2 | 0.59 ± 0.14 | 1.31 ± 0.25 | 1.24 ± 0.32 | 0.81 ± 0.21 |
|  |  | 15-keto PGF2alpha | 0.08 ± 0.03 | 0.18 ± 0.11 | 0.22 ± 0.11 | 0.14 ± 0.08 |
|  |  | 13,14dh-15k-PGE2 | ND | 0.34 ± 0.08 | ND | 0.19 ± 0.09 |
|  |  | tetranor PGEM | ND | ND | ND | ND |
|  |  | PGJ2 | 0.71 ± 0.11 | 0.47 ± 0.14 | 0.70 ± 0.16 | 0.86 ± 0.21 |
|  |  | D12-PGJ2 | ND | ND | ND | ND |
|  |  | 13,14dh-15k-PGD2 | ND | 0.36 ± 0.23 | ND | ND |
|  |  | 11dh-TXB2 | ND | ND | ND | ND |
|  |  | 2,3-dinor TXB2 | ND | ND | ND | ND |
|  |  | 13,14dh-15k-PGF2alpha | ND | ND | ND | ND |
|  |  | 8-isoPGF2alpha & 11bPGF2alpha | ND | ND | ND | ND |
|  |  | 19(R)-OH PGF2alpha & 20-OH PGF2alpha | ND | ND | ND | ND |
|  |  | Bicyclo PGE2 | ND | ND | ND | ND |
|  |  | 19(R)-OH PGE2 & 20-OH PGE2 | ND | ND | ND | ND |
|  |  | 15d-D12,14-PGJ2 | ND | ND | ND | ND |
|  |  | 6,15-diketo PGFalpha | ND | ND | ND | ND |
|  |  | iPF-VI | ND | ND | ND | ND |
|  | 20:5n-3 | TXB3 | ND | ND | ND | ND |
|  |  | 11dh TXB3 | ND | ND | ND | ND |
|  |  | PGD3 | ND | ND | ND | ND |
|  |  | PGE3 | ND | ND | ND | ND |
|  |  | PGF3alpha | ND | ND | ND | ND |
|  |  | 15d-D12,14-PGJ3 | ND | ND | ND | ND |

Values are mean ± SEM of 8-12 muscles from 4-6 rats per group. ND = Below limits of detection of the assay.

**Supplemental Tables 1F:** Lipoxygenase metabolite concentration (pg/mg) in the rat plantaris muscle in response to functional overload.

| Pathway | Substrate | Analyte | Synergist ablation time-point |  |  |  |
| --- | --- | --- | --- | --- | --- | --- |
|  |  |  | Control | 3D | 7D | 28D |
| Lipoxygenase | 18:2n-6 | 9-HODE | 10.20 ± 2.55 | 12.76 ± 3.62 | 9.36 ± 4.22 | 10.68 ± 2.52 |
|  |  | 9-OxoODE | 25.57 ± 4.54 | 29.34 ± 4.40 | 18.02 ± 2.83 | 42.75 ± 8.98 |
|  |  | 13-HODE | 86.61 ± 13.69 | 118.75 ± 25.86 | 69.53 ± 12.43 | 132.81 ± 26.42 |
|  |  | 13-OxoODE | 54.09 ± 10.69 | 72.70 ± 12.03 | 43.31 ± 6.13 | 85.30 ± 15.46 |
|  | 18:3n-3 | 9-HOTrE | 0.15 ± 0.07 | 0.70 ± 0.28 | 0.59 ± 0.17 | 0.77 ± 0.40 |
|  |  | 9-OxoOTrE | 0.97 ± 0.22 | 0.71 ± 0.16 | 0.53 ± 0.14 | 1.04 ± 0.21 |
|  |  | 13-HOTrE | 1.00 ± 0.27 | 2.39 ± 1.54 | 0.51 ± 0.24 | 0.97 ± 0.30 |
|  | 18:3n-6 | 13-HOTrE(gamma) | ND | ND | ND | ND |
|  | 20:2n-6 | 11-HEDE | 0.12 ± 0.05 | 0.13 ± 0.06 | 0.09 ± 0.04 | 0.29 ± 0.08 |
|  |  | 15-HEDE | ND | 0.42 ± 0.25 | 0.07 ± 0.03 | 0.23 ± 0.07 |
|  |  | 15-OxoEDE | ND | 0.06 ± 0.03 | ND | ND |
|  | 20:3n-9 | 5-HETrE | 0.15 ± 0.02 | 0.17 ± 0.03 | 0.11 ± 0.02 | 0.14 ± 0.03 |
|  |  | 8-HETrE | 0.15 ± 0.06 | 0.32 ± 0.08 | 0.21 ± 0.05 | 0.25 ± 0.07 |
|  | 20:4n-6 | 5-HETE | 4.86 ± 0.74 | 9.41 ± 1.18 | 5.68 ± 1.31 | 8.90 ± 1.92 |
|  |  | 5-oxoETE | 1.54 ± 0.29 | 2.71 ± 0.54 | 2.29 ± 0.40 | 2.83 ± 0.59 |
|  |  | 8-HETE | 2.83 ± 0.42 | 7.23 ± 1.79 | 3.34 ± 0.44 | 6.37 ± 1.33 |
|  |  | 9-HETE | 0.56 ± 0.24 | 1.20 ± 0.52 | 0.99 ± 0.46 | 1.30 ± 0.55 |
|  |  | 11-HETE | 9.41 ± 1.46 | 26.44 ± 6.30 | 11.84 ± 2.19 | 22.76 ± 4.37 |
|  |  | 12-HETE | 27.72 ± 7.44 | 270.16 ± 163.86 | 28.40 ± 10.25 | 46.81 ± 17.86 |
|  |  | 12-OxoETE | ND | ND | ND | ND |
|  |  | tetranor 12-HETE | 0.35 ± 0.08 | 1.19 ± 0.16 | 0.32 ± 0.06 | 0.33 ± 0.14 |
|  |  | 15-HETE | 7.48 ± 0.67 | 20.12 ± 6.17 | 8.67 ± 1.82 | 9.47 ± 1.44 |
|  |  | 15-OxoETE | 0.51 ± 0.13 | 0.66 ± 0.13 | 0.88 ± 0.19 | 0.46 ± 0.19 |
|  |  | 20-HETE | 12.98 ± 1.89 | 35.70 ± 4.76 | 24.76 ± 7.07 | 42.13 ± 7.07 |
|  |  | 5(S),12(S)-DiHETE | 0.45 ± 0.24 | 22.58 ± 13.56 | 0.96 ± 0.40 | 2.16 ± 0.68 |
|  |  | 5(S),15(S)-DiHETE | ND | 0.41 ± 0.27 | ND | ND |
|  |  | 8(S),15(S)-DiHETE | ND | ND | ND | ND |
|  |  | LTB4 | ND | ND | ND | ND |
|  |  | 5(S),6(S)-DiHETE | 0.23 ± 0.16 | 1.11 ± 0.73 | 0.14 ± 0.08 | 0.41 ± 0.31 |
|  |  | 5,6-DiHETE(n-3)) | ND | ND | ND | ND |
|  |  | 12-OxoLTB4 | ND | ND | ND | ND |
|  |  | 20-hydroxy LTB4 | ND | ND | ND | ND |
|  |  | 20-COOH LTB4 | ND | ND | ND | ND |
|  |  | 18-carboxy dinor LTB4 | ND | ND | ND | ND |
|  | 20:5n-3 | 5-HEPE | 0.63 ± 0.13 | 0.90 ± 0.22 | 0.78 ± 0.10 | 0.97 ± 0.18 |
|  |  | 8-HEPE | ND | ND | ND | ND |
|  |  | 9-HEPE | ND | ND | ND | ND |
|  |  | 11-HEPE | ND | 0.52 ± 0.28 | 0.24 ± 0.15 | ND |
|  |  | 12-HEPE | 3.27 ± 1.01 | 18.94 ± 8.20 | 2.30 ± 0.88 | 3.17 ± 1.07 |
|  |  | 15-HEPE | 0.24 ± 0.12 | 3.18 ± 2.93 | 0.44 ± 0.28 | 0.55 ± 0.31 |
|  |  | 18-HEPE | 1.32 ± 0.31 | 1.25 ± 0.26 | 0.92 ± 0.16 | 1.62 ± 0.30 |
|  |  | LTB5 | ND | ND | ND | ND |
|  | 22:6n-3 | 4-HDoHE | 4.69 ± 0.80 | 6.79 ± 1.10 | 3.67 ± 0.38 | 4.82 ± 0.74 |
|  |  | 7-HDoHE | 0.65 ± 0.17 | 1.02 ± 0.18 | 0.73 ± 0.25 | 0.90 ± 0.18 |
|  |  | 8-HDoHE | 1.92 ± 0.35 | 2.15 ± 0.69 | 1.53 ± 0.35 | 3.20 ± 0.76 |
|  |  | 10-HDoHE | 1.92 ± 0.47 | 4.24 ± 0.88 | 1.84 ± 0.34 | 2.62 ± 0.59 |
|  |  | 11-HDoHE | 1.49 ± 0.33 | 5.04 ± 1.69 | 1.19 ± 0.44 | 2.61 ± 0.45 |
|  |  | 13-HDoHE | 2.01 ± 0.27 | 3.97 ± 0.96 | 1.93 ± 0.32 | 3.08 ± 0.59 |
|  |  | 14-HDoHE | 3.77 ± 1.06 | 11.50 ± 3.23 | 3.31 ± 0.89 | 4.68 ± 1.44 |
|  |  | 16-HDoHE | 3.43 ± 0.55 | 5.48 ± 0.81 | 3.56 ± 0.87 | 3.64 ± 0.35 |
|  |  | 17-HDoHE | 1.20 ± 0.33 | 4.88 ± 2.20 | 1.08 ± 0.26 | 0.93 ± 0.20 |
|  |  | 20-HDoHE | 2.61 ± 0.46 | 3.33 ± 0.53 | 2.62 ± 0.31 | 3.20 ± 0.46 |

Values are mean ± SEM of 8-12 muscles from 4-6 rats per group. ND = Below limits of detection of the assay.

**Supplemental Tables 1G:**
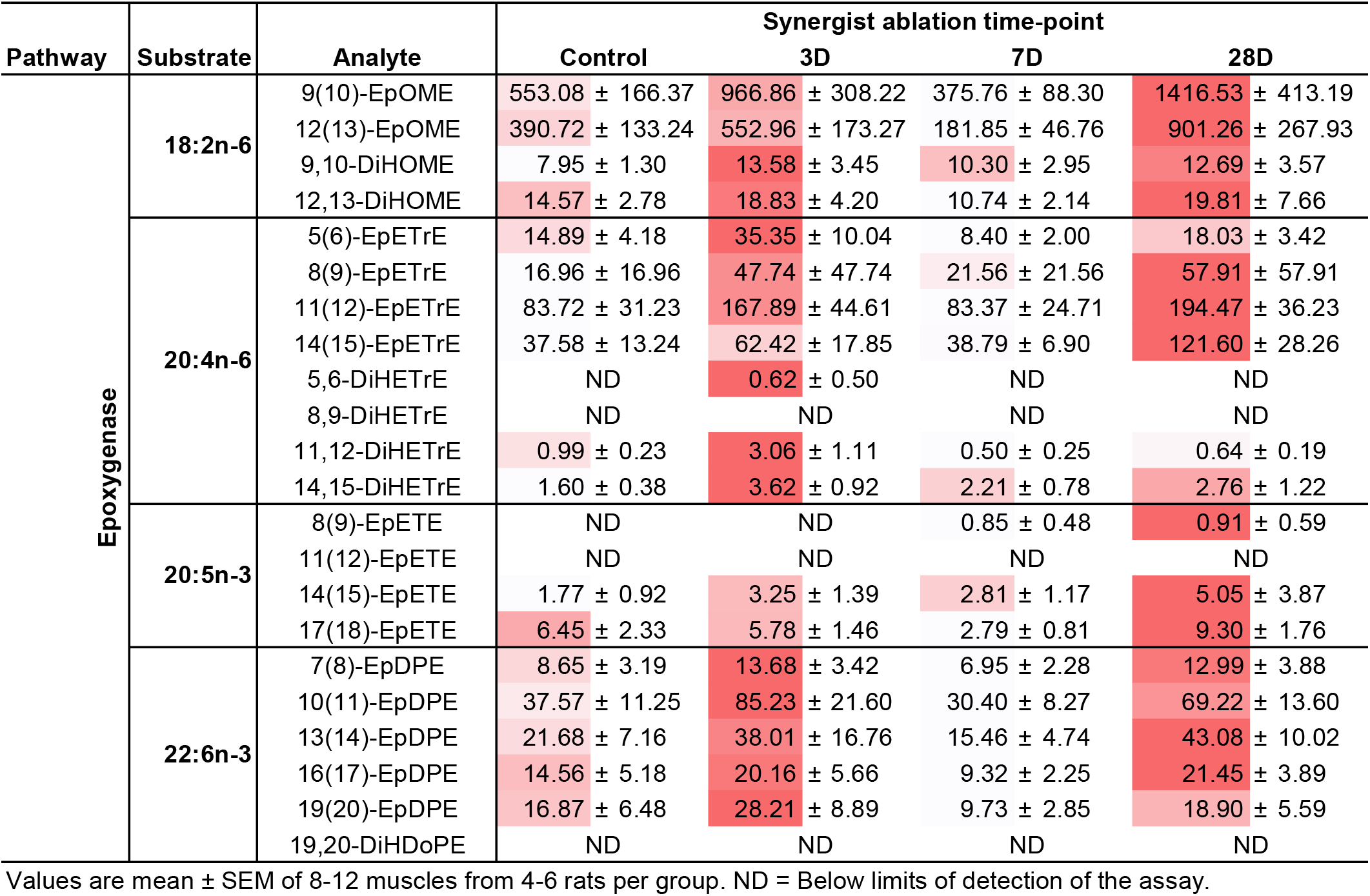
Epoxygenase metabolite concentration (pg/mg) in the rat plantaris muscle in response to functional overload.

| Pathway | Substrate | Analyte | Synergist ablation time-point |  |  |  |
| --- | --- | --- | --- | --- | --- | --- |
|  |  |  | Control | 3D | 7D | 28D |
| Epoxygenase | 18:2n-6 | 9(10)-EpOME | 553.08 ± 166.37 | 966.86 ± 308.22 | 375.76 ± 88.30 | 1416.53 ± 413.19 |
|  |  | 12(13)-EpOME | 390.72 ± 133.24 | 552.96 ± 173.27 | 181.85 ± 46.76 | 901.26 ± 267.93 |
|  |  | 9,10-DiHOME | 7.95 ± 1.30 | 13.58 ± 3.45 | 10.30 ± 2.95 | 12.69 ± 3.57 |
|  |  | 12,13-DiHOME | 14.57 ± 2.78 | 18.83 ± 4.20 | 10.74 ± 2.14 | 19.81 ± 7.66 |
|  | 20:4n-6 | 5(6)-EpETrE | 14.89 ± 4.18 | 35.35 ± 10.04 | 8.40 ± 2.00 | 18.03 ± 3.42 |
|  |  | 8(9)-EpETrE | 16.96 ± 16.96 | 47.74 ± 47.74 | 21.56 ± 21.56 | 57.91 ± 57.91 |
|  |  | 11(12)-EpETrE | 83.72 ± 31.23 | 167.89 ± 44.61 | 83.37 ± 24.71 | 194.47 ± 36.23 |
|  |  | 14(15)-EpETrE | 37.58 ± 13.24 | 62.42 ± 17.85 | 38.79 ± 6.90 | 121.60 ± 28.26 |
|  |  | 5,6-DiHETrE | ND | 0.62 ± 0.50 | ND | ND |
|  |  | 8,9-DiHETrE | ND | ND | ND | ND |
|  |  | 11,12-DiHETrE | 0.99 ± 0.23 | 3.06 ± 1.11 | 0.50 ± 0.25 | 0.64 ± 0.19 |
|  |  | 14,15-DiHETrE | 1.60 ± 0.38 | 3.62 ± 0.92 | 2.21 ± 0.78 | 2.76 ± 1.22 |
|  | 20:5n-3 | 8(9)-EpETE | ND | ND | 0.85 ± 0.48 | 0.91 ± 0.59 |
|  |  | 11(12)-EpETE | ND | ND | ND | ND |
|  |  | 14(15)-EpETE | 1.77 ± 0.92 | 3.25 ± 1.39 | 2.81 ± 1.17 | 5.05 ± 3.87 |
|  |  | 17(18)-EpETE | 6.45 ± 2.33 | 5.78 ± 1.46 | 2.79 ± 0.81 | 9.30 ± 1.76 |
|  | 22:6n-3 | 7(8)-EpDPE | 8.65 ± 3.19 | 13.68 ± 3.42 | 6.95 ± 2.28 | 12.99 ± 3.88 |
|  |  | 10(11)-EpDPE | 37.57 ± 11.25 | 85.23 ± 21.60 | 30.40 ± 8.27 | 69.22 ± 13.60 |
|  |  | 13(14)-EpDPE | 21.68 ± 7.16 | 38.01 ± 16.76 | 15.46 ± 4.74 | 43.08 ± 10.02 |
|  |  | 16(17)-EpDPE | 14.56 ± 5.18 | 20.16 ± 5.66 | 9.32 ± 2.25 | 21.45 ± 3.89 |
|  |  | 19(20)-EpDPE | 16.87 ± 6.48 | 28.21 ± 8.89 | 9.73 ± 2.85 | 18.90 ± 5.59 |
|  |  | 19,20-DiHDoPE | ND | ND | ND | ND |

Values are mean ± SEM of 8-12 muscles from 4-6 rats per group. ND = Below limits of detection of the assay.

**Supplemental Tables 1H:**
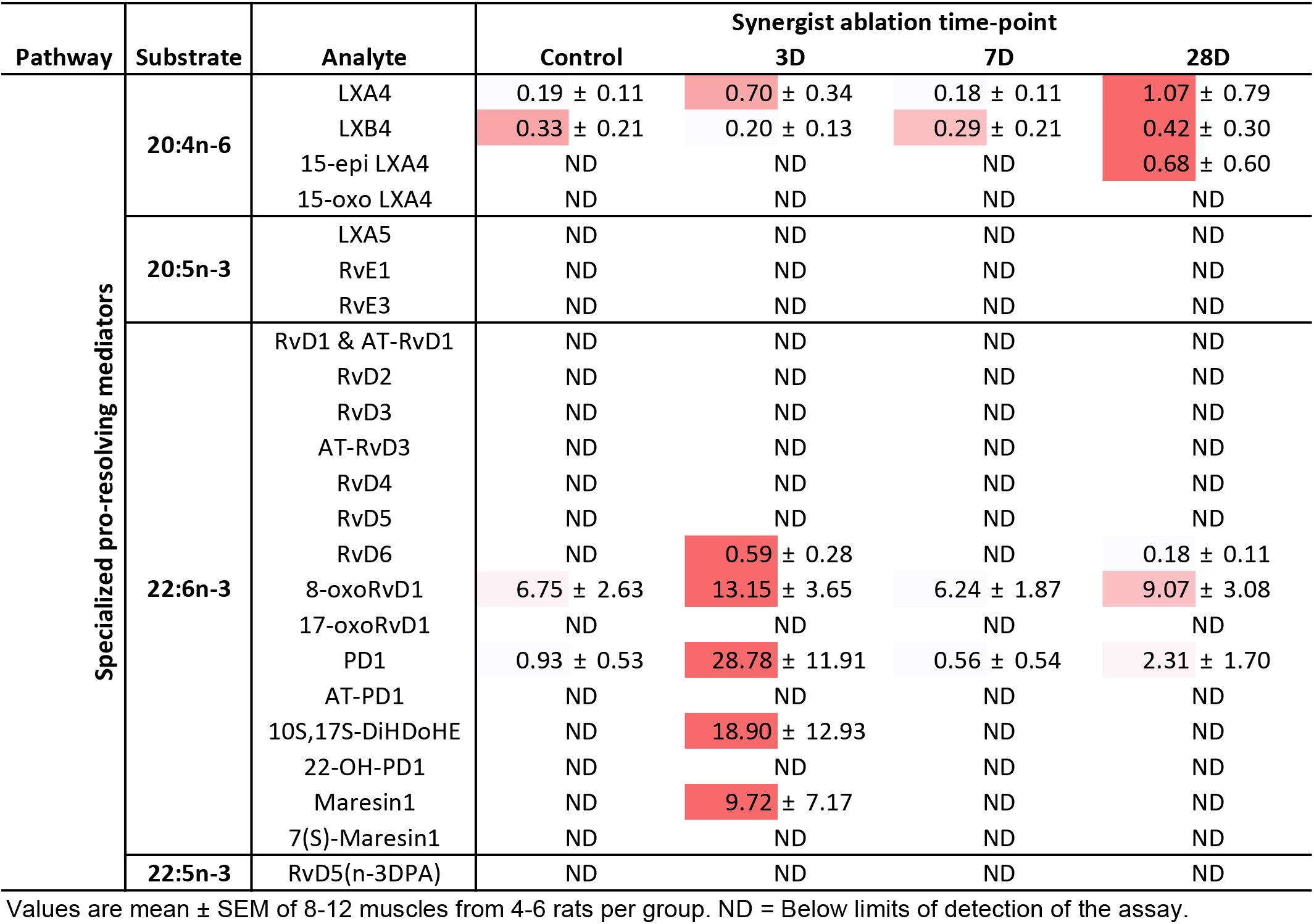
Specialized pro-resolving mediator concentration (pg/ mg) in the rat plantaris muscle in response to functional overload.

